# Glutarate regulates T cell function and metabolism

**DOI:** 10.1101/2022.10.20.513065

**Authors:** Eleanor Minogue, Pedro P. Cunha, Alessandro Quaranta, Javier Zurita, Shiv Sah Teli, Brennan J. Wadsworth, Rob Hughes, Guinevere L. Grice, Pedro Velica, David Bargiela, Laura Barbieri, Craig E. Wheelock, James A. Nathan, Peppi Koivunen, Iosifina P. Foskolou, Randall S. Johnson

**Affiliations:** Department of Physiology, Development and Neuroscience, University of Cambridge, UK; Department of Cell and Molecular Biology, Karolinska Institutet, Sweden; Unit of Integrative Metabolomics, Institute of Environmental Medicine, Karolinska Institutet, Department of Respiratory Medicine and Allergy, Karolinska University Hospital, Stockholm, Sweden; Biocenter Oulu, Faculty of Biochemistry and Molecular Medicine, Oulu Centre for Cell-Matrix Research, University of Oulu, Finland; Cambridge Institute of Therapeutic Immunology & Infectious Disease (CITIID), Jeffrey Cheah Biomedical Centre, Cambridge Biomedical Campus, University of Cambridge, Cambridge, CB2

## Abstract

T cell function is influenced by several metabolites; some acting through enzymatic inhibition of α-KG-dependent dioxygenases (αKGDDs), others, through post-translational modification of lysines in important targets. We show here that glutarate, a product of amino acid catabolism, has the capacity to do both, with effects on T cell function and differentiation. Glutarate exerts those effects through αKGDD inhibition and through direct regulation of T cell metabolism via post-translational modification of the pyruvate dehydrogenase E2 subunit. Diethyl-glutarate, a cell-permeable form of glutarate, alters CD8^+^ T cell differentiation and increases cytotoxicity against target cells. *In vivo* administration of the compound reduces tumor growth and is correlated with increased levels of both peripheral and intratumoral cytotoxic CD8^+^ T cells. These results demonstrate that glutarate regulates both T cell metabolism and differentiation, with a potential role in the improvement of T cell immunotherapy.

## Introduction

T cell activation results in quiescent naïve T cells differentiating into rapidly proliferating effector cells. This transition involves dynamic metabolic reprograming in which cells increase consumption of glucose, amino acids, and fatty acids to meet energy demands (Blagih *et al*., 2015; Carr *et al*., 2010; Frauwirth *et al*., 2002; Nakaya *et al*., 2014; Sinclair *et al*., 2013). Following an immune response, some T cells remain and transition to memory cells, which are primed for rapid recall responses, while others become exhausted and die by apoptosis (Kaech and Cui, 2012; Sallusto *et al*., 2010). T cell differentiation, memory cell formation, and survival are regulated by several intrinsic factors, including transcriptional and epigenetic regulation (Crompton *et al*., 2016; Harland *et al*., 2014; Kaech and Cui, 2012; Russ *et al*., 2014). These processes are also regulated by extrinsic factors, including nutrient and oxygen availability (Caldwell *et al*., 2001; Kedia-Mehta and Finlay, 2019; Palazon *et al*., 2017; Ross *et al*., 2021; Wei *et al*., 2017).

Some metabolites are important modulators of T cell differentiation and function. In recent years, our laboratory and others have illustrated the importance of the S enantiomer of 2-hydroxyglutarate (S-2HG) in T cell fate (Bunse *et al*., 2018; Tyrakis *et al*., 2016). We have shown that supplementation of T cell cultures with esterified forms of S-2HG enhances T cell immunotherapy in preclinical models by promoting the development of central memory T (T_CM_) cells (Foskolou *et al*., 2020; Tyrakis *et al*., 2016).

2HG, as well as succinate and fumarate, are structural analogues of alpha-ketoglutarate (αKG) and can act as competitive inhibitors of αKG-dependent dioxygenases (αKGDDs) (Chowdhury *et al*., 2011; Koivunen *et al*., 2007, 2012; Laukka *et al*., 2016; Xiao *et al*., 2012). There are currently over 60 identified αKGDDs, and many of these enzymes play pivotal roles in cellular function (Baksh and Finley, 2020; Losman *et al*., 2020; Morris *et al*., 2019). It has been shown that S-2HG exerts its T cell modulatory effects via competitive inhibition of many of the αKGDDs (Tyrakis *et al*., 2016). We sought to determine whether molecules which are structurally similar to S-2HG could exert similar effects on T cell differentiation and function.

Here we describe our finding that glutarate is a potent inhibitor of several αKGDDs and has an unexpected and important role in cellular metabolism and T cell differentiation. Glutarate can cause shifts in the regulation of the hypoxia inducible factor (HIF), and alterations in histone and DNA methylation. We further demonstrate that glutarate can modify the pyruvate dehydrogenase E2 subunit (PDHE2) post-translationally, altering its activity, and thus act as a novel regulator of pyruvate metabolism. These wide-ranging functions for glutarate indicate a central role for the metabolite as a major regulator of metabolism and immune function.

## Results

### Diethyl Glutarate (DEG) is a regulator of T cell differentiation

We first sought to investigate if structurally similar compounds to 2HG could increase the T_CM_ population of CD8^+^ T cells, acting similarly to the S enantiomer of 2HG. We identified 19 structural analogues of 2HG; 8 were commercially available, and 11 were synthesized de-novo for the purposes of this study (Figure S1A). We used esterified versions of most of the compounds employed so as to increase potency and aid intracellular translocation. The octyl-ester form of S-2HG was used as a positive control. We screened the compounds for effects on T cell differentiation by making them 400 μM (the optimal S-2HG dose in CD8^+^ T cells (Tyrakis *et al*., 2016)) in each culture media during activation of isolated primary murine CD8^+^ T cells. After seven days of culture, we assessed the abundance of CD62L^hi^/CD44^hi^, T_CM_-like T cells by flow cytometry (Figure 1A). We additionally determined the effect of each compound on cell growth and viability (Figure 1B and Figure 1SB). Of the 19 test compounds, only diethyl-glutarate (DEG) significantly increased the abundance of CD62L^hi^/CD44^hi^ CD8^+^ T cells in this assay (Figure 1A). Further, treatment with DEG had no negative effects on either cell viability or proliferation (Figure 1B and Figure S1B).

**Figure 1:**
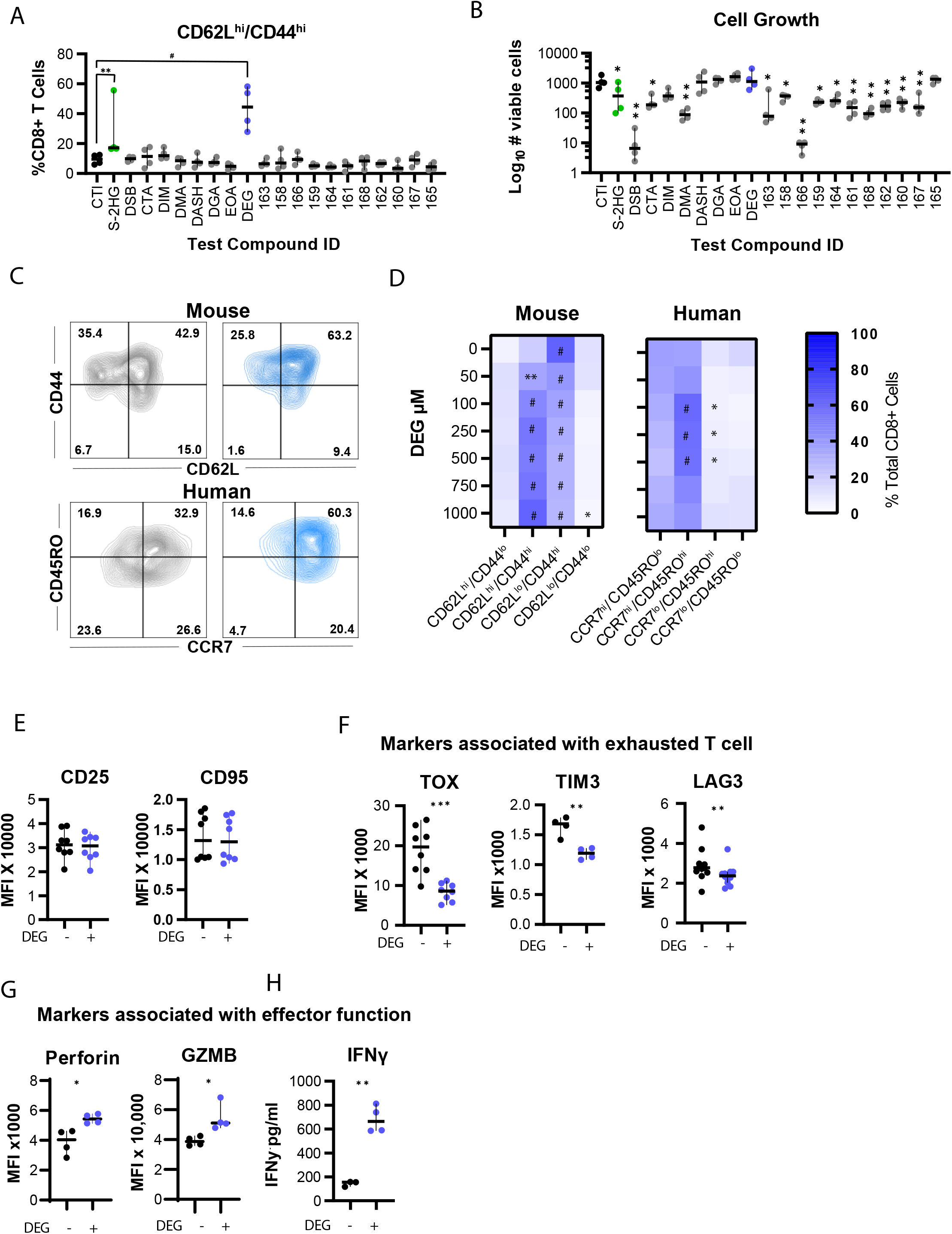
Diethyl Glutarate (DEG) is a regulator of T cell differentiation. (A and B) Percentage of CD62L^hi^/CD44^hi^ and number of total murine CD8^+^ T cells following 7 days of treatment with 400μM of test compound. Ordinary one-way ANOVA relative to control treated cells (CT); n=4. (C) Representative flow cytometry counter plots representing the expression of CD44/CD62L in mouse CD8^+^ T cells and CD45RO/CCR7 in human CD8^+^ T cells treated +/− 500μM DEG for 7 days. Pre-gated on live, single, CD8^+^ events. (D) Heatmap representing proportion of mouse CD8^+^ T cells expressing CD62L/CD44 (left) and the proportion of human CD8^+^ T cells expressing CCR7/CD45RO (right) following 7 days of treatment with increasing concentrations of DEG. Two-way ANOVA relative to CT; n=4-5. (E) Cell surface expression of CD25 and CD95 on human CD8^+^ T cells following 7 days of treatment +/− DEG 500μM, as determined by flow cytometry and shown by Median Fluorescence Intensity (MFI). Paired test; n=8. (F) Expression of markers associated with exhaustion on human CD8^+^ T cells following 7-10 days of treatment +/− DEG 500μM, as determined by flow cytometry and shown by MFI. Paired t test; n= 4-8. (G) Expression of cytotoxic granules on human CD8^+^ T cells following 7 days of treatment +/− DEG 500μM, as determined by flow cytometry and shown by MFI. Paired t test; n=4. (H) IFNγ secretion by human CD8^+^ T cells, 3 days post activation and treatment +/− DEG 500μM, as measured by ELISA. Paired t-test; n=4 All scatter plots show median and 95% confidence interval (CI). Each dot represents one donor (human or murine as indicated). *p<0.05, **p<0.01, ***p<0.001, #P<0.0001.

We then performed titration experiments and found that DEG increased the CD62L^hi^/CD44^hi^ T_CM_ population in a dose-dependent manner, at concentrations similar to effective doses of octyl-ester S-2HG (Figure S1C). DEG did not display any negative effects on cell growth, even at 1 mM concentrations, in contrary to octyl-ester S-2HG, which decreases cell growth in the highest concentrations used (Figure S1D). Further analysis of memory populations in both mouse and human DEG-treated CD8^+^ T cells revealed that after 7 days of culture, DEG increased the T_CM_ population, as represented by CD62L^hi^/CD44^hi^ in mouse and CCR7^hi^/CD45RO^hi^ in human CD8^+^ T cells; and decreased the effector memory T cell (T_EM_) population as represented by CD62L^lo^/CD44^hi^ in mouse and CCR7^lo^/CD45RO^hi^ in human CD8^+^ T cells (Figure 1C-D).

Following stimulation with anti-CD3/CD28 microbeads, DEG treated CD8^+^ T cells became fully activated, as determined by the cell surface expression of the markers CD25 and CD95 (Figure 1E). Following T cell activation, naïve T cells differentiate into effector cells. Effector T cells can become exhausted, and die shortly thereafter by apoptosis, or become long-lived memory cells. Following our observation above that DEG alters memory T cell formation rates, we examined multiple exhaustion-associated markers in treated cells, and found that TOX, a transcription factor known to orchestrate exhaustion (Khan *et al*., 2019; Sekine *et al*., 2020), and the exhaustion markers TIM3 and LAG3, were all significantly downregulated in human CD8^+^ T cell cultures following 10 days of DEG treatment (Figure 1F).

The primary function of CD8^+^ T cells is elimination of target cells, which is mediated by exocytosis of lysosomes containing cytotoxic proteins such as granzyme b and perforin and the secretion of cytokines such as IFN-γ. We found that DEG treatment increased expression of the perforin and granzyme B proteins, and the secretion of IFNγ (Figure 1G-H).

Together, this data illustrates the DEG can alter CD8^+^ T cell differentiation and results in the formation of more effector-like CD8^+^ T cells.

### Glutarate is a novel T cell immunometabolite

DEG is a di-esterified form of the metabolite glutaric acid (glutarate). We next wished to confirm that T cells are capable of cleaving the di-esters in DEG, and that glutarate levels thus rise in DEG treated cells. To do this, we performed isotope tracing of ^13^C_5_-labelled DEG in human CD8^+^ T cells. We found that DEG is rapidly converted into glutarate following cellular uptake (Figure S2A-B). The data shown here indicate that this processing of diethyl glutarate into glutarate occurs within minutes after DEG is added to T cell culture media. Consistent with this, intracellular DEG could not be detected intracellularly; this was seen as soon as 15 minutes after DEG treatment (Figure S2B). This indicates a complete conversion of DEG to glutarate occurs almost immediately after DEG enters T cells.

Glutarate, in its coenzyme A (CoA) form, is an intermediate of both tryptophan and lysine catabolism (Borsook *et al*., 1948; Gholson *et al*., 1959) (Figure 2A). Glutaryl-CoA can either be further broken down to crotonyl-CoA by the enzyme glutaryl-CoA dehydrogenase (GCDH), or form glutarate (Figure 2A) (Dwyer *et al*., 2000). The glutarate to glutaryl-CoA reaction is mediated by Succinyl-CoA:Glutamate-CoA transferase (SUGCT) (Marlaire *et al*., 2013). The ratio of crotonyl-CoA to glutarate production is highly regulated, and excessive glutarate accumulation is cytotoxic (Goodman *et al*., 1975). Excess glutarate formed due to loss of function mutations in GCDH gives rise to the metabolic disorder Glutaric Aciduria Type 1, which can lead to developmental issues in children and, ultimately, acute metabolic crises leading to fatal seizures (Goodman *et al*., 1975; Kölker *et al*., 2000; Kyllerman *et al*., 2004; Strauss *et al*., 2003).

**Figure 2:**
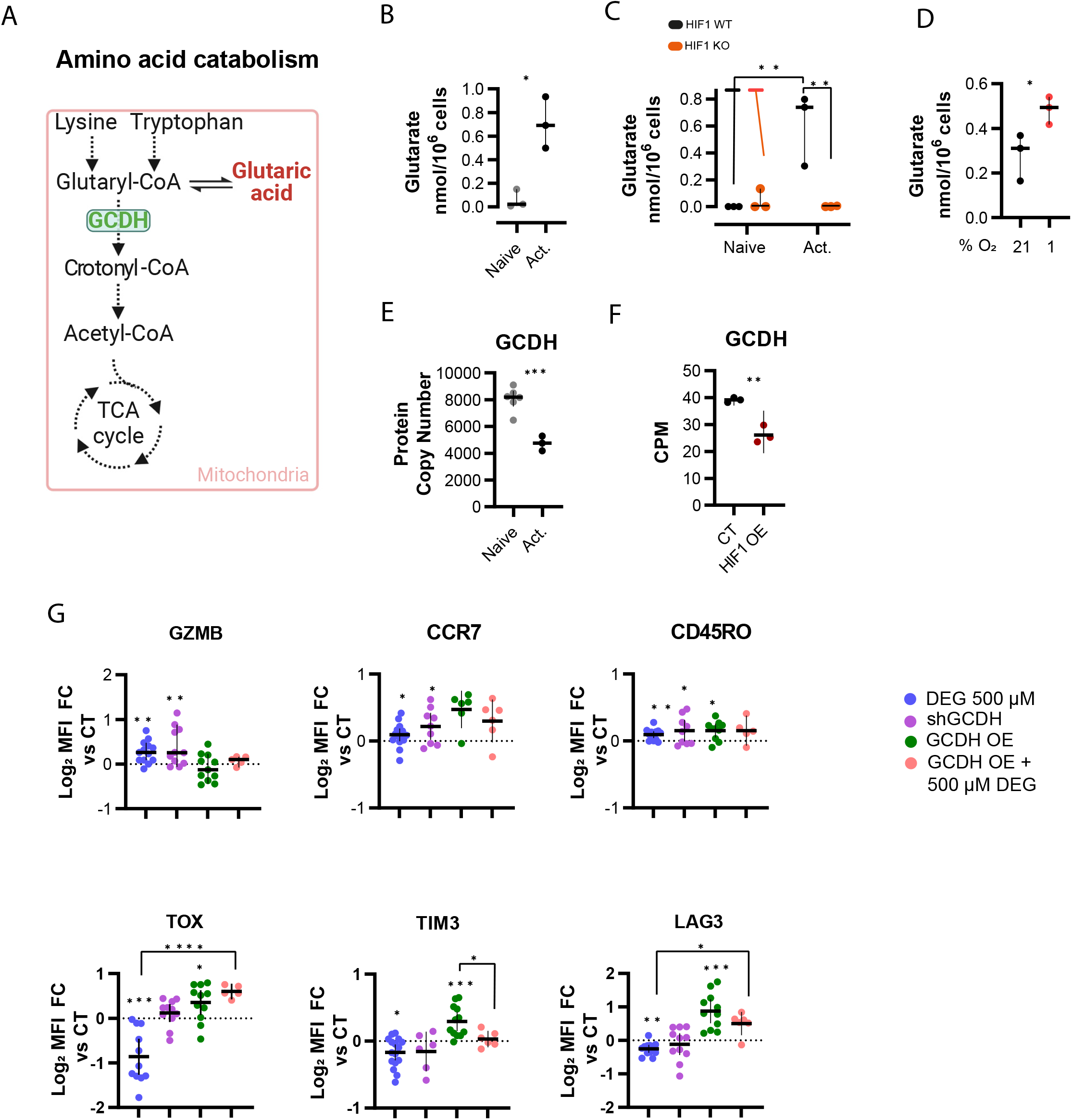
Glutarate is a novel T cell metabolite with immunomodulatory properties. (A) Pathway of lysine and tryptophan catabolism. (B) Glutarate levels in naïve and 72 hr activated mouse CD8^+^ T cells. Paired t test; n=3. (C) Glutarate levels in naïve or 72 hr activated CD8^+^ T cells isolated from Hif-1αfl/fl or Hif-1αfl/fl dlckcre mice (HIF ko). Two-way ANOVA; n=3. (D) Glutarate levels in mouse CD8^+^ T cells cultured at 21% or 1% oxygen for 24 h from day 5 post activation. Paired t test; n=3. (E) GCDH protein copy number in naïve or activated CD8^+^ T cells from P14 transgenic mice. Unaired t test; n=3. Data from ImmPRes database (Howden *et al*., 2019). (F) GCDH counts per million (CPM) in HIF overexpressing (HIF OE) mouse CD8^+^ T cells. Unpaired t test; n=3. (Veliça *et al*., 2021). (G) Expression of markers of T cell differentiation and exhaustion in human CD8^+^ T cells following 7-10 days treatment +/− DEG 500μM and/or transduction with either shGCDH or GCDH OE vector. Expression as determined by flow cytometry and shown as MFI FC relative to relative untreated control (black dotted line). One sample t test; n=6-12. (J All scatter plot shows median + 95% CI where each dot represents one donor (murine or human as indicated). *p<0.05, **p<0.01, ***p<0.001.

Although glutarate is a product of amino acid catabolism and excessive glutarate accumulation can be pathological, there is a very limited literature describing the role of glutarate in physiological settings, and essentially no work describing the role of glutarate in immune cells. We thus sought to determine the actual concentrations of glutarate in naïve and activated CD8^+^ T cells. We used mass spectrometry to determine intracellular levels of endogenous glutarate in T cells and found that levels are substantially increased following T cell activation (Figure 2B). Glutarate levels in activated T cells were significantly higher than the levels of the metabolite 2HG, a known immunometabolite (Figure S1C). Using previously published CD8+ T cell volumes, we estimate that the concentration of glutarate in activated CD8+ T cells is approximately 0.6 mM. Treating activated CD8^+^ T cells with 500 μM DEG increased intracellular glutarate concentrations by approximately 6-fold (relative uptake of labelled glutarate shown in Figure S2D).

It is known that T cell metabolism is greatly altered after T cell activation, and a major factor driving these metabolic changes is the hypoxia inducible factor 1a (HIF-1α) (Nakamura *et al*., 2005). Thus, we asked if the observed increase of glutarate levels after T cell activation was HIF1 dependent. We used HIF1 proficient and HIF1 null (HIF1KO) CD8^+^ T cells and found that the activation-induced increase in cellular glutarate seen in HIF1 proficient cells is essentially lost in HIF1KO cells, indicating a role for the HIF1 transcription factor in modulating T cell glutarate levels (Figure 2C). We also found that culturing T cells at 1% oxygen for 24 hours increased glutarate levels approximately 2-fold compared to culturing T cells at 21% oxygen levels (Figure 2D).

As previously alluded to, glutarate levels are in part determined by the activity of the GCDH enzyme. To determine if the increased glutarate levels we observed were due to decreased levels of GCDH expression, we used published datasets (Howden *et al*., 2019; Veliça *et al*., 2021) and found that GCDH protein copy number is decreased following T cell activation (Figure 2E). In addition, GCDH gene expression levels decrease in HIF-1α-overexpressing, but not HIF-2α-overexpressing T cells (Figure 2F and Figure S2E).

Following the observation that GCDH levels fluctuate in CD8^+^ T cells and that this is correlated with intracellular glutarate levels, we next created genetically modified GCDH constructs to determine the role of endogenously generated glutarate in CD8^+^ T cells. We wished in this way to either increase intracellular glutarate levels, via silencing of GCDH (shGCDH); or decrease intracellular glutarate levels, via overexpression of GCDH (GCDH_OE) (Figure S2F). CD8^+^ T cells treated with DEG and CD8^+^ T cells expressing shGCDH showed similar increases in expression of granzyme b, CCR7 and CD45RO (Figure 2G). CD8^+^ T cells over-expressing GCDH had similar levels of granzyme b and CCR7 relative to their respective controls but, as seen with DEG treated cells, they had increased expression of CD45RO (Figure 2G). Strikingly, GCDH over-expression had the opposite effect of DEG treatment on the expression of the exhaustion markers TOX, TIM3 and LAG3: DEG supplementation decreases the expression of TOX, TIM3 and LAG3, while GCDH over-expression increases the levels of each of those markers significantly (Figure 2G). Supplementing GCDH overexpression CD8^+^ T cells with DEG was able to rescue the effects of GCDH overexpression on TIM3 and LAG3 (Figure 2G), indicating that high levels of GCDH can be offset by increasing glutarate levels via DEG supplementation.

### Glutarate is an inhibitor of αKGDDs

Given the structural similarities to 2HG, fumarate and succinate, all competitive inhibitors of αKGDDs (Chowdhury *et al*., 2011; Koivunen *et al*., 2007, 2012; Laukka *et al*., 2016; Xiao *et al*., 2012), we postulated that glutarate may exert its effects by inhibiting αKG-dependent enzymes. To determine this, we focused on three subfamilies of the most intensively studied αKGDDs: the DNA demethylating Ten-eleven translocation methylcytosine dioxygenases (TETs), the HIF prolyl 4-hydroxylases (HIF-P4Hs), and the histone lysine demethylases (KDMs); all of whose reactions are depicted in Figure S3A. In cell free enzymatic activity assays, glutarate was found to inhibit recombinant TET2 with an IC50 of 2 mM (Figure 3A). This inhibition of TET2 correlated with an observed decrease in the TET2 reaction product 5hmC in a dose-dependent manner in human CD8^+^ T cells cultured with DEG (Figure 3B). This indicates that glutarate is capable of inhibiting TET2 activity and thus inhibiting DNA demethylation.

**Figure 3:**
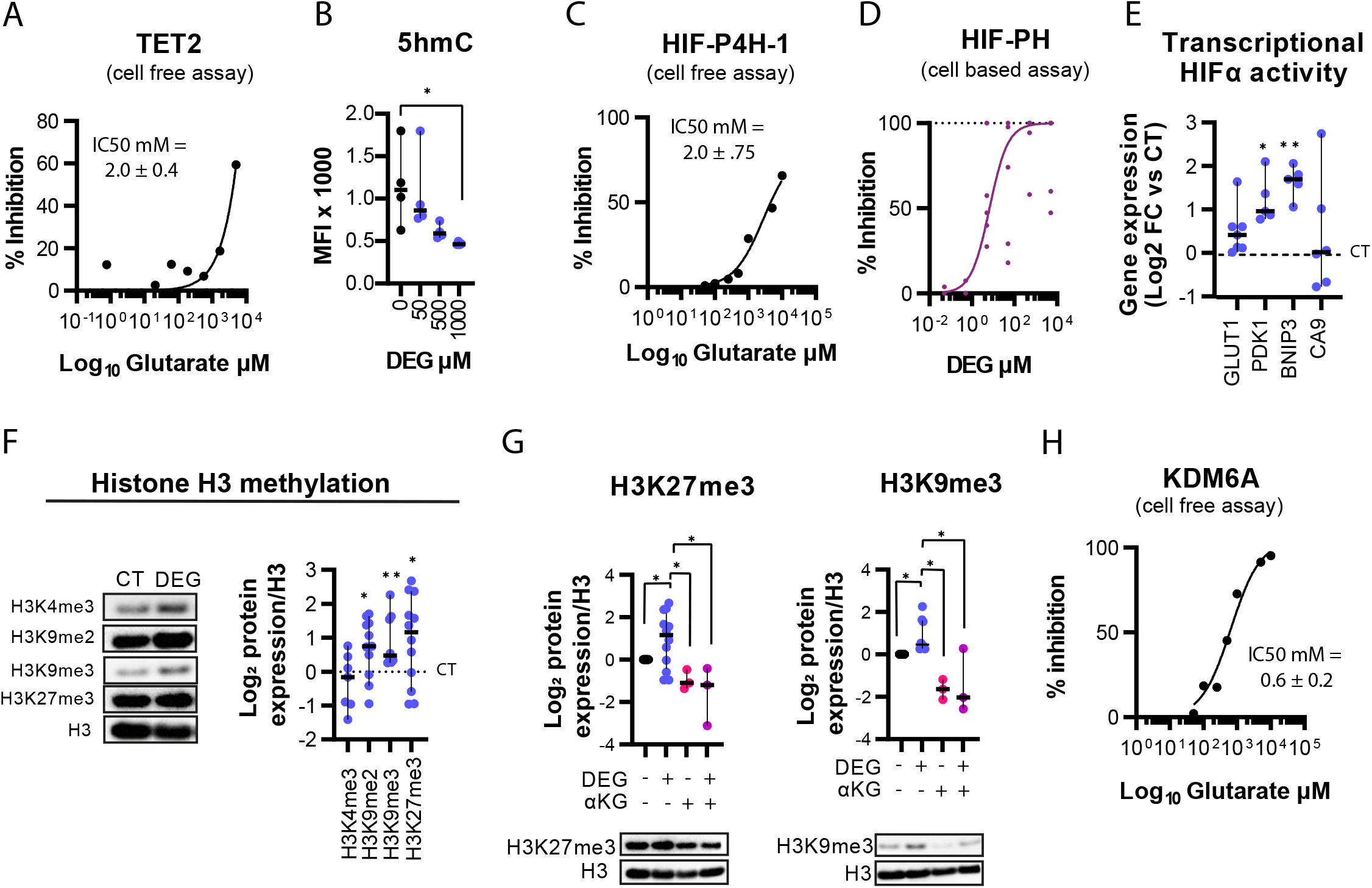
Glutarate is an inhibitor of αKG-dependent reactions. (A) Cell free enzymatic inhibition assay for TET2. Inhibition was determined using increasing concentrations of glutarate. Graph shows Michaelis-Menten line of best fit. (B) 5hmC MFI, as determined by flow cytometry, in human CD8^+^ T cells following 7 days of treatment with increasing concentrations of DEG. Ordinary one-way ANOVA; n=4. (C) Cell free enzymatic inhibition assay for HIF-P4-H1. Values are means of at least 3 independent experiments. Graph shows Michaelis-Menten line of best fit. (D) HIF-PH activity luciferase reporter assay in MEFs treated for 16h with DEG. N=4-6. (E) qPCR analysis of CD8^+^ T cells treated with DEG for 7 days. Each dot represents one human donor relative to untreated donor control (dotted black line), normalised to HPRT. One sample t test; n=5-8. (F) Representative western blot and log_2_ FC in protein expression of specific histone H3 (H3) methylation sites in human CD8^+^ T cells treated for 7 days +/− DEG 500μM. Each dot represents one human donor normalised to total H3, and relative to untreated control (dotted black line). One-sample Wilcoxon test; n=6-12. (G) Representative western blots and log2 FC protein expression of H3K27me3 and H3K9me3 in human CD8^+^ T cells treated with +/− DEG and/or αKG for 7 days. Kruskal-Wallis test; n=3-12. (H) Cell free enzymatic inhibition assay for KDM6A. Values are means of at least 3 independent experiments. Graph shows Michaelis-Menten line of best fit. All scatter plots show median + 95% CI, where each dot represents one human donor. *p<0.05, **p<0.01.

Glutarate was also found to inhibit the HIF-P4H-1 (HIF -proyl4hydroxylase-1) enzyme *in vitro*, with an IC50 of 2 mM in a cell free assay (Figure 3C). To test HIF-P4H inhibition via DEG treatment in a cellular system, we employed a luciferase reporter assay driven by a Gal4 response element (GRE-luc), pFLAG-Gal4-mHIF-1α NTAD (Pereira *et al*., 2003). Cells were concurrently transfected with ampGRE-luc and a plasmid including constitutive expression of a fusion protein linking a FLAG-tagged Gal4 DNA-binding domain to one of the transcription activation domains of murine HIF-1α. The N-terminal transcription activation domain (NTAD) contains proline residues that are hydroxylated by HIF-P4H enzymes to target the protein for degradation, thus luciferase expression is controlled by HIF-P4H hydroxylation activity against the pFLAG-Gal4-mHIF-1α NTAD. We found that DEG can inhibit HIF-P4H activity in this cellular system (Figure 3D). This inhibition of HIF-P4H was correlated with increased expression of multiple HIF target genes in DEG treated CD8^+^ T cells (Figure 3E).

To determine if glutarate also inhibits KDM enzymes, we cultured CD8^+^ T cells with DEG for 7 days and determined the methylation status of a number of histone H3 (H3) lysine residues (Figure 3F). DEG treatment of CD8^+^ T cells increased di- and tri-methylation of H3K9 and trimethylation of H3K27 (Figure 3F). This H3K9 and H3K27 trimethylation was abrogated by the addition of equimolar amounts of αKG, further suggesting that DEG is a competitive inhibitor of αKGDDs with regards to this enzymatic reaction (Figure 3G). H3K9me2 levels were not reduced with the addition of αKG (Figure S3B). As KDM6A is responsible for the demethylation of H3K27, we performed a cell free enzymatic activity assay with this enzyme and found that glutarate inhibited KDM6A with an IC50 of 0.6 mM (Figure 3H).

Here we show that glutarate is an inhibitor of specific αKGDDs, namely, the TET2, HIF-P4H, and KDM6A enzymes. This inhibition correlates with multiple changes to cellular genetic and epigenetic profiles and is correlated with increased expression of specific HIF targets, increased histone methylation, and increased DNA demethylation.

### Glutarylation of Pyruvate Dehydrogenase E2

Some metabolites can give rise to post-translational modifications (PTMs) on specific residues in proteins (Chen *et al*., 2007; Kim *et al*., 2006; Weinert *et al*., 2013). Glutarylation, a recently discovered PTM, uses glutaryl-CoA as a substrate (Figure 4A) and can occur both non-enzymatically (Tan *et al*., 2014) and enzymatically (Bao *et al*., 2019) on lysine residues. Deglutarylation is reported to be mediated by SIRT5 (Tan *et al*., 2014). It has recently been shown in studies of mouse liver and brain that glutarylation can modify a range of proteins (Schmiesing *et al*., 2018; Tan *et al*., 2014). Given that glutarate levels increase after CD8^+^ T cell activation (Figure 2B-C), we questioned if glutarylation can also occur in CD8^+^ T cells.

**Figure 4:**
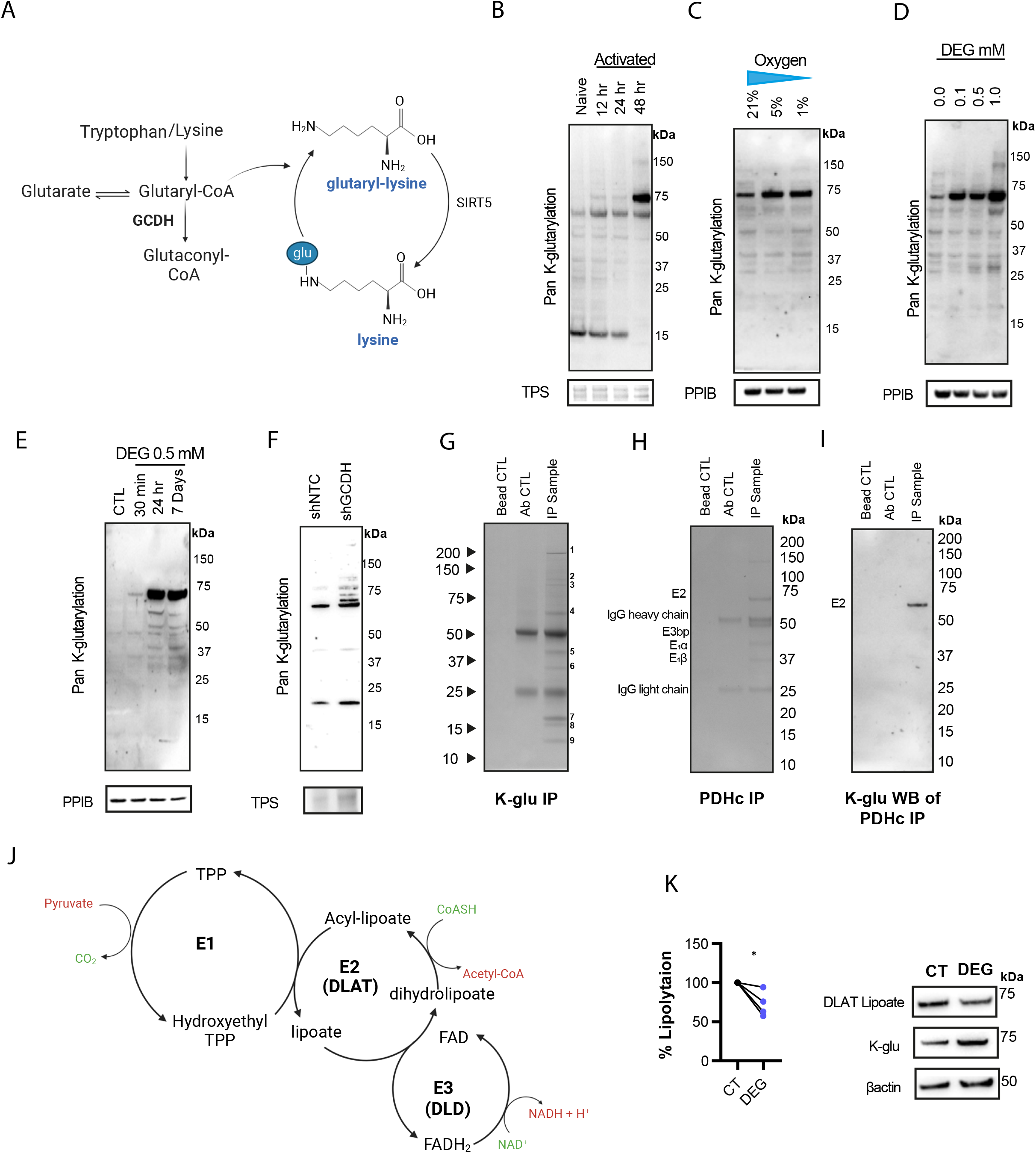
Glutarylation of PDHE2. (A) Model of protein lysine-glutarylation. (B) Representative western of naïve and activated human CD8^+^ T cells. N=3. (C) Representative western blot of activated human CD8^+^ T cells cultured at different oxygen tensions (21%, 5% or 1%). N=4. (D) Representative western of human CD8^+^ T cells cultured with increasing concentrations of DEG for 7 days. N=4. (E) Representative western of activated human CD8^+^ T cells cultured with DEG for various time lengths as indicated. All samples were harvested 7 days post activation. N=4. (F) Representative western of CD8^+^ T cells transduced with shNTC or shGCDH vectors. RQR8^+^ cells were purified by FACS prior to analysis. N=3. (G) Coomassie staining of proteins immunoprecipitated with a pan-k-glutarylation antibody and separated by SDA-PAGE. 30×10^6^ mouse CD8^+^ T cells, 7 days post activation were used. (H) PDHc isolated from 30×10^6^ mouse CD8^+^ T cells by immunoprecipitation. (I) k-glutarylation WB of immunoprecipitated PDHc as described in Figure 4H. (J) Mode of PDHc. (K) Lipoate and PDHE2 glutarylation levels in HeLa cells treated with DEG. (K) % lipoylation of DLAT (PDHE2) in HEK293 cells treated with DEG for 24h as determined by western blot analysis and relative to βactin. Unpaired t test; n=4. *p<0.05.

We first performed whole protein lysate western blot analysis with a pan-lysine-glutarylation (K-glu) antibody (Tan *et al*., 2014; Xie *et al*., 2021) and found that glutarylation is clearly detectable in CD8^+^ T cells (Figure 4B). We also found that glutarylation patterns change strikingly upon CD8^+^ T cell activation (Figure 4B). These glutarylation patterns were also altered when cells were exposed to different oxygen tensions (Figure 4C and Figure S4A), and when treated with DEG (Figure 4D and Figure S4B). Interestingly, shifts in glutarylation could be detected following just 30 minutes of culture with DEG (Figure 4E and Figure S4C) and these were increased with partial GCDH silencing (Figure 4F). As shown in Figure 2, DEG is converted within 15 minutes to glutarate intracellularly; indicating that the DEG-induced glutarylation observed after 30 minutes of DEG treatment is likely due to increases in intracellular glutarate.

We observed only a small number of proteins, detectable by glutarylation western blots, whose glutarylation changed markedly during T cell activation (Figure 4B-F). We were particularly interested in determining the identity of a protein of approximately 70kDa, which by total protein western blot analysis appeared to be the most glutarylated protein in T cells after activation. This was also the protein most markedly affected by both hypoxia and by DEG exposure (Figure 4B-E and Figure S4A-C).

In order to establish the identity of the glutarylated proteins in CD8^+^ T cells, we performed an immunoprecipitation assay with a pan-lysine-glutarylation (K-glu) antibody (Figure 4G). We excised the observed bands and used mass spectrometry in conjunction with the SwissProt protein database to identify them (Figure S4D). Proteins with the highest PEP score in each band are listed in Figure S4E.

Three non-cytoskeletal/histone proteins were positively identified; (1) 2-oxoglutarate dehydrogenase (OGDH), (2) dihydrolipoyl lysine-residue acetyltransferase (DLAT) (also known as the E2 subunit of the pyruvate dehydrogenase complex - PDHE2), and (3) pyruvate dehydrogenase E1 component subunit beta (PDHE1β) (Figure S4E). Glutarylation targets were confirmed by reverse immunoprecipitation, in which an immunoprecipitation with the target antibody was performed and then the resulting western blot was probed with the K-glu antibody. Immunoprecipitation and subsequent western blot analysis with this K-glu antibody of the PDH complex (PDHc) (which includes PDHE2 and PDHE1β) revealed that it was the PDHE2 subunit alone that is clearly and detectably glutarylated (Figure 4 H-I). Immunoprecipitation and subsequent western blot analysis with the K-glu antibody revealed that OGDH is not detectably glutarylated (Figure S4F-G). Thus, the PDHE2 subunit (67kDa) was the only glutarylation target we could both identify by mass spectrometry and subsequently confirm by reverse immunoprecipitation.

The PDHc catalyses the conversion of pyruvate to acetyl-CoA, and thus links glycolysis and the TCA cycle. PDHc is comprised of multiple copies of three catalytic enzymes: pyruvate dehydrogenase (E1), dihydrolipoamide acetyltransferase (DLAT) (PDHE2), and dihydrolipoamide dehydrogenase (DLD) (E3) (Figure 4J). The E2 subunit is the functional core of the PDHc and provides the catalytic activity by cyclical reduction and oxidation of its lipoyl domains, channelling substrates between in individual enzymes’ active sites in the PDHc. The multidomain E2 subunit consists of the C-terminal catalytic domain, the peripheral subunit-binding domain and the N-terminal end of 1-3 lipoyl domains (Bleile *et al*., 1979; Green *et al*., 1992; Perham *et al*., 1981; Stephens *et al*., 1983). The lipoyl domains contain one lipoic acid covalently attached to a lysine residue (Packman *et al*., 1991). Following the observation that lysine residues of the E2 subunit of the PDHc can be glutarylated, we sought to determine if lysine glutarylation was disrupting lipoic acid conjugation to E2 (DLAT) lysine residues and thereby disrupting normal PDHc functioning. We treated HEK cells with DEG and observed reduced DLAT E2 lipoate in the presence of lysine glutarylation, indicating that lysine glutarylation of E2 inhibits normal lipoate function (Figure 4K).

These data illustrate that glutarylation occurs in CD8^+^ T cells, the glutarylation status changes after activation and identifies PDHE2 as a glutarylation target, whilst providing evidence in HEK cells that glutarylation disrupts normal PDHc lipoylation, potentially altering catalytic activity of the complex.

### Glutarate modulates mitochondrial function by inhibiting pyruvate dehydrogenase via both glutarylation and phosphorylation

PDHc is a key player in the regulation of metabolism, and links glycolysis to the TCA cycle (Figure 5A) (Patel *et al*., 2014).. Following our observation that the E2 subunit can be glutarylated and that this results in disruption of the E2 lipoyl domains, we hypothesised that glutarylation would decrease PDHc enzymatic activity, resulting in increased glycolysis.

**Figure 5:**
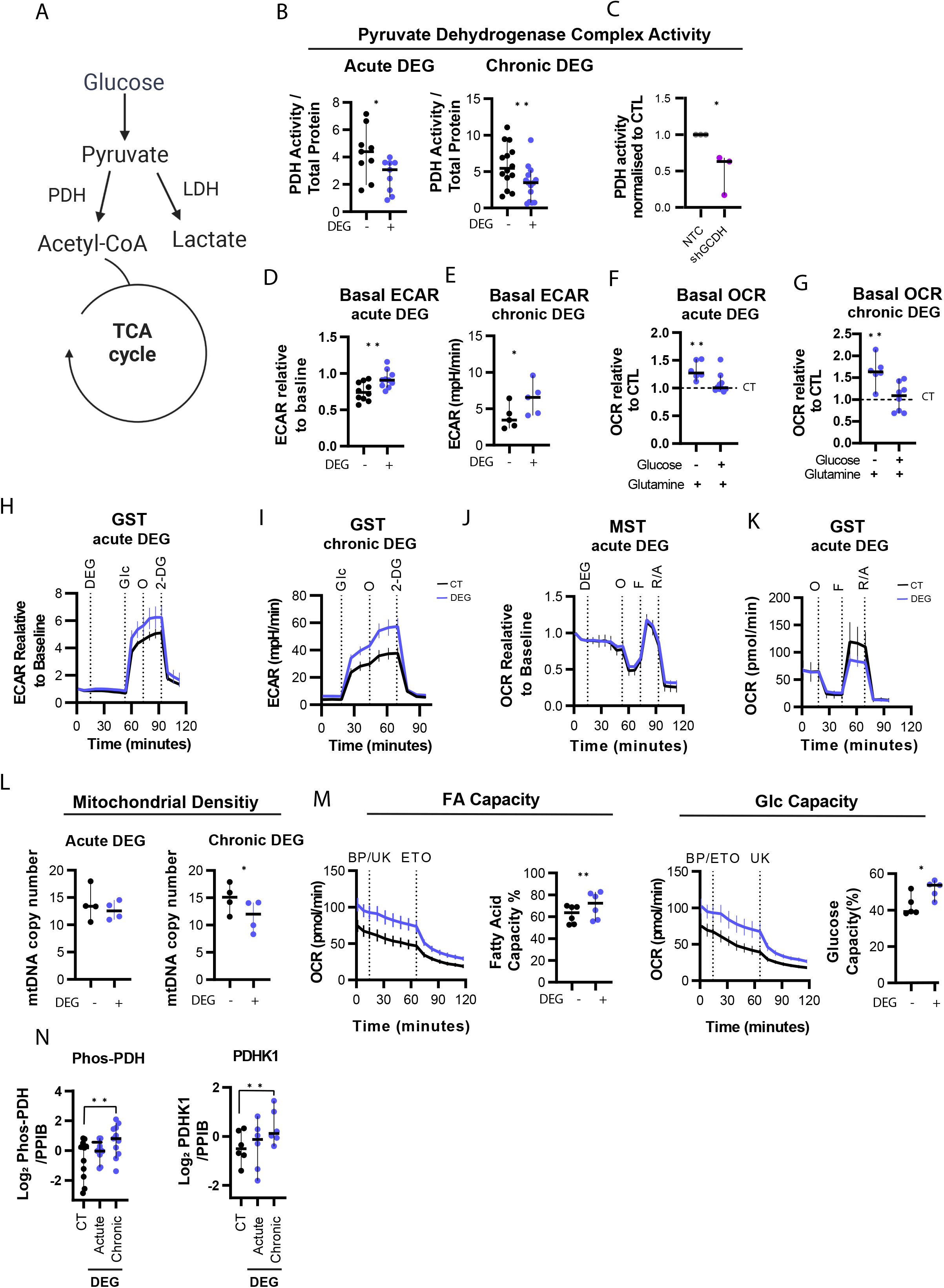
Glutarate modulates metabolism via both glutarylation and phosphorylation of the PDHE2. (A) Schematic of glycolysis and TCA cycle. (B) PDHc activity of human CD8^+^ T cells treated with DEG for 30 min (acute) or 7 days (chronic), normalised to total protein concentration. Paired t test; n=9-14. (C) PDHc activity of CD8^+^ T cells transduced with shGCDH relative to NTC control. RQR8^+^ cells were purified by FACS before assay. Unpaired t test; n= 3. (D) Basal ECAR levels in CD8^+^ T cells as determined by Seahorse FX analysis following 30 min DEG treatment via seahorse injection. ECAR normalised to ECAR prior to DEG/CT injection. Paired t test; 30 min n=10. (E) Basal ECAR levels in CD8^+^ T cells following 7 days of treatment +/− DEG 500μM as determined by Seahorse FX analysis Paired t test; 30 min n=5. (F) Basal OCR levels in CD8^+^ T cells as determined by Seahorse FX analysis following 30 min DEG treatment via seahorse injection. During assay cells were plated in FX media +/− glucose and glutamine as indicated. OCR represented at FC relative to untreated control. One-sample t test; n=6-10. (G) Basal OCR levels in CD8^+^ T cells following 7 days of treatment +/− DEG 500μM as determined by Seahorse FX analysis. During assay cells were plates in FX media +/− glucose and glutamine as indicated. One sample t test; n=6-10. (H) Seahorse XF Analysis of ECAR levels in CD8^+^ T cells (7 days post activation). Injections 1-4: DEG/CT, Glucose (Glc), Oligomycin (O), 2-Deoxy-D-Glucose (2-DG), as indicated. Mean and Error with 95% CI; n=10. (I) ECAR measurements during a standard GST of CD8^+^ T cells treated with +/− DEG 500μM for 7 days. Mean and Error with 95% CI; n=6. (J) Seahorse XF Analysis of OCR levels in CD8^+^ T cells (7 days post activation). Injections 1-4: DEG/CT, Oligomycin (O), Carbonyl cyanide 4- (trifluoromethoxy)phenylhydrazone (F), Rotenone/Antimycin (R/A). Mean and Error with 95% CI; n=10. (K) ECAR measurements during a standard GST of CD8^+^ T cells treated with +/− DEG 500μM for 7 days. Mean and Error with 95% CI; n=6. (L) mtDNA copy number in cells treated with +/− DEG 500μM. Acute treatment of CD8^+^ T cells 7 days post activation for 24h. Chronic treatment of CD8^+^ T cells for 7 days, from activation. Paired t test; n=4 (M) Mitochondrial fatty acid (FA) and glucose (Glc) oxidative capacity of CD8^+^ T cells treated +/− DEG 500μM for 7 days, as determined by OCR measurement pre and post addition of BPTES (BP), UK5099 (UK) and etomoxir (ETO) as indicated in Figure S5C. Paired t test; n=6. (N) Protein expression of phosphorylated PDH (ser239) and PDHK1, as determined by western blot analysis and normalised to PPIB. Acute treatment of CD8^+^ T cells 7 days post activation for 30 min. Chronic treatment of CD8^+^ T cells for 7 days, from activation. Protein levels normalised to PPIB or complex V expression as indicate. Wilcoxon test; acute n=10, chronic n=5. All scatter plots show median + 95% CI where each dot represents one human. *p<0.05, **p<0.01.

To test this, we performed a direct enzymatic activity assay of the PDHc on CD8^+^ T cell lysates, and found that both acute (30 minute) and chronic (7 day) exposure to DEG results in decreased PDHc activity (Figure 5B). Reduced PDHc activity was also observed in T cells with partially silenced GCDH (Figure 5C). As would be expected, this reduction in PDHc activity resulted in increases in the basal extracellular acidification rate (ECAR) (Figure 5D-E). Surprisingly, basal oxygen consumption rate (OCR) was also increased in CD8^+^ T cells treated with DEG, after acute and chronic treatment, when assayed in the absence of glucose. OCR levels were unchanged when glucose was present (Figure 5F-G).

To explore further the metabolic consequences of the observed glutarate induced reduction in PDHc activity we performed both a glycolysis stress test (GST) and a mitochondrial stress test (MST) in CD8^+^ T cells. Acute effects were determined by treating CD8^+^ T cells with DEG for 30 minutes whilst plated in the Seahorse XY Analyser, prior to preforming a standard GST or MST. Chronic effects were determined by preformed standard GST and MST analysis in CD8^+^ T cells that had been treated with DEG *in vitro* for 7 days prior to assay. Both acute and chronic exposure to DEG increased glycolysis (Figure 5H-I and Figure S5A). Acute exposure to DEG did not alter mitochondrial oxidation during the MST (Figure 5J and Figure S5B). However, following chronic DEG exposure, CD8^+^ T cells had a reduced maximal OCR following treatment with the uncoupling agent, FCCP (Figure 5K and Figure S5B).

Following the observation that chronic, but not acute, DEG treatment results in reduced maximal achievable OCR, we sought to determine if chronic DEG treatment alters mitochondrial density. Quantification of mitochondrial DNA (mtDNA) copy number revealed that chronic, but not acute, DEG treatment of CD8^+^ T cells results in a significant reduction in mtDNA copy number (Figure 5L). Given that basal OCR was not negatively affected but was in fact increased when glucose was limited (Figure 5G), we sought to determine alternative fuel pathways that chronically treated CD8^+^ T cells may be utilising following DEG treatment. Again, using the Seahorse FX Analyser, we used different combinations of inhibitors of glutamine, glucose, and fatty acid mitochondrial oxidation (BPTES, UK5099 and etomoxir respectively) and determined their effect on oxygen consumption (Figure S5C). This analysis revealed that chronic DEG treatment increased CD8^+^ T cell mitochondrial fatty acid and glucose oxidation (Figure 5M). We did not observe a change in glutamine oxidative capacity following DEG treatment (Figure S5D). The observed increase in capacity for fatty acid oxidation was correlated with the upregulation of genes associated with fatty acid metabolism, including, fatty acid synthase (FASN), fatty acid binding protein 2 (FABP2) and carnitine palmitoyl transferase 1A (CPT1A) (Figure S5E) in DEG treated T cells.

PDHc activity is tightly regulated by substrate availability and by phosphorylation by pyruvate dehydrogenase kinase (PDHK). To determine if the alterations observed were caused solely by glutarylation, we quantified PDHc protein levels, phosphorylation levels and levels of PDHK in CD8^+^ T cells treated with DEG. Acute DEG treatment does not alter any of these parameters significantly (Figure 5N and Figure S5F), providing further evidence that the changes in PDHc activity and glycolysis observed with acute DEG treatment are due to glutarylation and subsequent disruption of lipoylation of PDHE2.

Chronic DEG treatment of CD8^+^ T cells does not reduce PHDc total protein levels (taking overall changes in mitochondrial density observed with chronic DEG treatment into consideration, by normalising to complex V protein expression) (Figure S5F). Both expression of pyruvate dehydrogenase kinase-1 (PDHK1) and phosphorylation of PDHc are increased with chronic DEG treatment (Figure 5N and S5G) which is consistent with PDHK1 being a direct HIF target, as shown in Figure 3.

These data illustrate that glutarate can modulate CD8^+^ T cells metabolism via glutarylation and phosphorylation of the PDHc. Glutarylation of PDHE2 occurs rapidly after glutarate exposure and results in rapid reductions in PDHc activity and in glycolysis. Chronic glutarate exposure alters cellular metabolism via both glutarylation and phosphorylation of PDHc; with the latter likely induced via HIF-P4H-1 inhibition, as glutarate acts as a competitive inhibitor of that enzyme.

### Glutarate reduces tumor growth, and increases T cell numbers in vivo and enhances tumor infiltration

Metabolic profiles of CD8^+^ T cells are known to drive effector functions (Bevilacqua *et al*., 2022). Given the observed increases in glycolysis described above, in addition to the increases in secretion of the cytotoxic granules, granzyme b and perforin seen following DEG treatment of CD8^+^ T cells, we postulated that they may also have increased cytotoxic function.

To test this, after 7 days of culture in DEG containing medium, antigen specific cytotoxicity was determined in both a human CAR-T cell system derived from human donor CD8^+^ T cells, and a murine system using transgenic OT-1 CD8^+^ T cells (Figure 6A). Human CD8^+^ T cells were activated for 24h and then transduced with a CD19-CAR-T cell vector and cultured with or without DEG. After 7 days of culture, CAR-T cells were co-cultured with CD19^+^ Raji cells and 16 hours later, supernatant was collected to determine the amount of T cell IFN-γ secretion by ELISA; cytotoxicity was determined by quantifying the live CD19^+^ Raji cells remaining by flow cytometry (Figure 6B). DEG treated human CD19-CAR-T cells had both increased IFNγ production and showed an increase in killing of CD19^+^ Raji target cells in this assay, when compared to untreated control CD19-CAR-T cells (Figure 6B).

**Figure 6:**
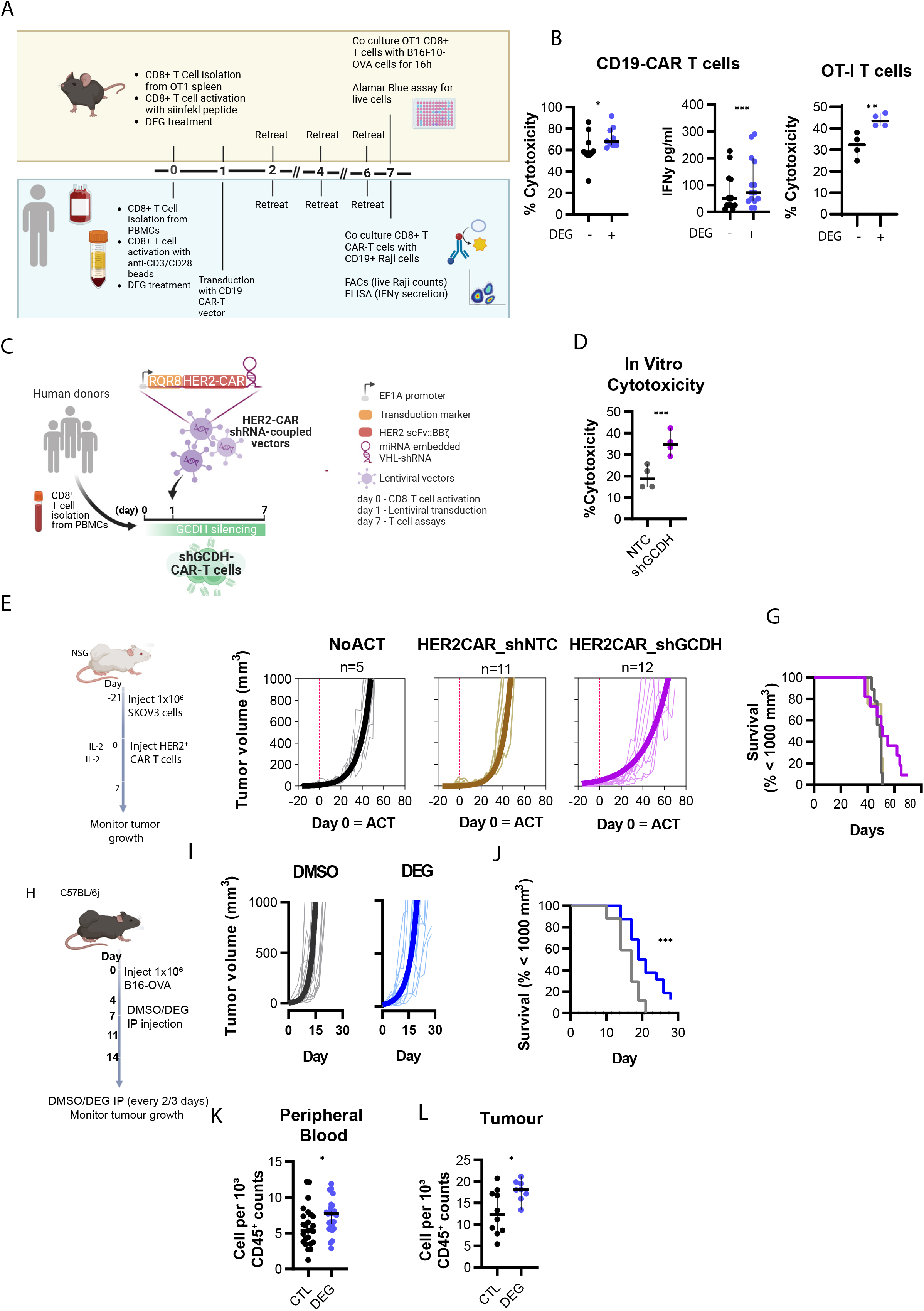
Glutarate reduces tumor growth, and increases T cell numbers and tumor infiltration. (A) Model of *in vitro* cytotoxicity assays. (B) Cytotoxicity and IFNγ expression (left) of CD19-CAR T cells following co culture with CD19^+^ Raji cells. Cytotoxicity of OT-1 following co culture with B16F10-OVA cells (right). CAR-T/OT-1 T cells were treated +/− DEG 500μM for 7 days prior to assay. Paired t test; n=9-12. (C) Model of HER2-CAR-T_shRNA cell generation. (D) Percentage cytotoxicity of HER2-CAR T cells with embedded shGCDH or NTC as determined by alamar blue assay following co-culture with HER2 expressing SKOV3 cells. Paired t test; n=4. (E) Adoptive cell therapy (ACT) model with CAR-T cells. 1×10^6^ SKOV3 cells were implanted subcutaneously (s.c.) in NSG animals that were injected 21 days later with human shNTC- or shGCDH-expressing CAR-T cells generated as shown in Figure 6C. Tumour growth was monitored every 2-3 days until day 70. (F) Tumor growth data. Thin lines represent tumor growth from individual animals and thick lines represent an exponential (Malthusian) growth curve. No ATC n=5, HER2-CAR-T_shNTC n=11, HER2-CAR-T_shGCDH n=12. (G) Survival curves using 1000mm^3^ in tumor volume as the threshold. Log-rank (mantel-Cox) test. (H) Tumor growth model. 1.0×10^6^ B16F10 cells were s.c. injected in C57BL/6j mice. From day 4 post tumor inoculation, mice were injected intraperitoneally with 10mg/kg DEG, or control, every 2-3 days. On day 14 after tumor inoculation, peripheral blood, tumor, spleen and tumor draining lymph node from some animals were processed to single cell suspensions and analysed by flow cytometry. Tumor growth was monitored until day 30. (I) Tumor growth data. Thin lines represent tumor growth from individual animals and thick lines represent an exponential (Malthusian) growth curve (n=16). (J) Survival curves using 1000mm^3^ in tumor volume as the threshold. Log-rank (mantel-Cox) test; n=14. (K) Frequency of CD8^+^ T cells in the peripheral blood, 14 days post tumor inoculation. Mann-Whitney test; n=25. (L) Frequency of CD8^+^ T cells in the tumours. Mann-Whitney test; n=14-16. All scatter plots show median and 95% CI where each dot represents one donor (human or murine as indicated). *p<0.05, **p<0.01, ***p<0.001.

To test if DEG increases cytotoxicity in a transgenic TCR murine system, OT-1 CD8^+^ T cells were used in conjunction with B16F10 melanoma tumor cells expressing the OVA peptide recognised by the OT-1 transgenic TCR (B16F10-OVA) (Figure 6A). DEG treated OT-1 CD8^+^ T cells killed significantly more B16F10-OVA cells when compared to untreated OT-1 cells (Figure 6B). Culture of human CD19-CAR-T cells or murine OT-1 T cells with DEG for 7 days did not affect cell growth (Figure S6A-B).

To investigate if endogenously increased glutarate levels in CD8^+^ T cells can also increase cytotoxic efficacy of CD8^+^ T cells, we created a human HER2-CAR-T system containing an miRNA-embedded shRNA against GCDH (HER2-CAR-T_shGCDH) (Figure 6C). HER2-CAR-T cells with a non-targeted control (NTC) (HER2-CAR-T_shNTC) were used as a control. CD8^+^ T cells expressing the HER2-CAR-T_shGCDH had significantly increased cytotoxicity against target HER2^+^ SKOV3 ovarian cancer cells compared to control HER2-CAR-T_shNTC cells (Figure 6D). Cell growth was not affected in the shGCDH-expressing cells (Figure S6C).

While CAR-T cell therapy has shown considerable clinical efficacy in the treatment of B cell lymphomas and leukemia, the successful treatment of solid tumors has been hindered in part due to the inhospitable tumor micro-environment (TME) (Marofi *et al*., 2021; Sterner and Sterner, 2021). Given our observations that increased intracellular glutarate, via both DEG treatment and partial GCDH silencing, results in more glycolytic cells that secrete higher levels of cytotoxic granules, whilst also increasing *in vitro* cytotoxicity, we performed *in vivo* tumor growth experiments in which mice with HER2^+^ SKOV3 tumors were treated with HER2-CAR-T_shGCDH cells (Figure 6E). Tumors in mice treated with HER2-CAR-T_shGCDH cells grew more slowly than tumors in mice treated with control HER2-CAR-T_NTC cells (Figure 6F). However, we did not observe a difference in the overall survival of the mice (Figure 6G).

We then questioned whether direct administration of DEG to tumor-bearing mice would result in a slowing of tumor growth. DEG is non-toxic and is approved for use as a food additive (Dionisio *et al*., 2018). To determine whether it might be able to be used therapeutically, we carried out an experiment in which mice with B16F10 melanoma tumors were injected with 10mg/kg DEG intraperitoneally every 48 to 72 hours (Figure 6H). This treatment regimen significantly slowed tumor growth and increased survival of the DEG-treated mice (Figure 6I-J). To investigate if the observed effect was due to a cytotoxic effect of DEG in tumour cells, we cultured tumour cells with various levels of DEG. We did not observe any cytotoxicity or cell growth reductions when DEG was added to a number of different tumor cell lines, including B16F10 tumor cells (Figure S6D-E), indicating that DEG does not generally affect tumor cell growth.

To determine the mechanism underlying the reduced tumor growth *in vivo*, we again undertook treatment of B16F10 tumor bearing mice with 10mg/kg i.p. DEG injection or control i.p. injection every 48 to 72 hours, as described above. Ten days after the start of DEG treatment, we harvested peripheral blood, tumor, spleen and tumor draining lymph nodes from the experimental animals. Single cell suspensions were generated from each sample and the quantity of several immune cell populations was determined (Figure S6F). CD3^+^ T cells, CD4^+^ T cells, CD8^+^ T cells, B cells, neutrophils, NK cells, monocytes, dendritic cells (DCs) and macrophages were assayed for, and this analysis revealed a significant increase in both blood CD8^+^ T cells and tumor infiltrated CD8^+^ T cells in the DEG treated animals relative to controls (Figure 6K-L). No changes in other immune populations examined were observed, apart from a slight decrease in circulating B cells (Figure S6G).

These data illustrate that glutarate can enhance CD8+ T cell cytotoxicity and that *in vivo* treatment of tumor bearing mice with esterified glutarate can reduce tumor growth and increase animal survival, highlighting glutarate as a potential cancer therapy.

## Discussion

In this study we identify the immunomodulatory capacity of the metabolite glutarate whilst also providing evidence that glutarate can influence cellular metabolic regulation by acting as both an inhibitor of αKGDDs, and by direct post-translational modification of the PHDc.

The intrinsic link between metabolism and cellular function was first highlighted by the discovery of mutations in genes encoding metabolic enzymes. Mutations in fumarate hydratase, succinate dehydrogenase, and isocitrate dehydrogenase (IDH), all of which are key enzymes of the TCA cycle, lead to the accumulation of fumarate, succinate and 2HG respectively. It is well established that at high levels, each of these metabolites can act as disease-driving signalling molecules. However, these are also essential metabolites in the regulation of many cellular functions across a range of cell types (Bourgeron *et al*., 1995; Parsons *et al*., 2008; Ryan *et al*., 2019; Tomlinson *et al*., 2002). Moreover, in recent years it has become clear that each of these metabolites play pivotal roles in immune cell regulation and in the proper orchestration of immune response (Arts *et al*., 2016; Jha *et al*., 2015; Tannahill *et al*., 2013; Tyrakis *et al*., 2016; Xu *et al*., 2017).

Mutations in the GCDH gene lead to chronically high levels of glutarate, which can be detrimental to normal development (Goodman *et al*., 1975). There is also some evidence that mutations in GCDH, like mutations in IDH, can lead to tumor development (Russi *et al*., 2018). To date GCDH and glutarate have almost exclusively been studied in the context of the disease syndrome of Glutaric Aciduria Type 1, stemming from inborn GCDH mutations. However, we illustrate here that glutarate can play a central role in regulation of cell physiology, differentiation, and metabolism. By acting as an inhibitor of αKGDDs, glutarate alters the transcriptional and epigenetic landscape of cells, via inhibition of HIF-P4H and KDMs/TETs respectively. We use CD8^+^ T cells as a model system here, but note that glutarate has this inhibitory property both in cell free assays, and in a range of other cell types; thus these effects are likely not exclusive to CD8^+^ T cells.

In an immune cell context, it is known that loss of TET2 expression promotes differentiation of CD8^+^ T cells into T_CM_ cells (Carty *et al*., 2018) and that a disrupted TET2 gene can promote therapeutic efficacy of CAR-T cells (Fraietta *et al*., 2018). We have illustrated here that glutarate treatment of CD8^+^ T cells results in direct TET2 inhibition, in turn resulting in increased levels of T_CM_ populations in activated CD8^+^ T cells and correlated with increased killing of target cells. It is also well established that cytotoxic CD8^+^ T cells require high glycolytic rates to sustain their function (Buck *et al*., 2015). We show here that following HIF-P4H-1 inhibition by glutarate, transcription of HIF target genes, including the glycolytic genes GLUT1 and PDHK1, is induced; this is correlated with increased glycolysis and in increased effector function of these cells.

There are over 60 αKGDDs, thus the effects observed in glutarate treated cells and in cells in which GCDH has been genetically manipulated are likely to be a complex interplay between many cellular reactions. It is also important to note here that the high levels of glutarate observed in patients with Glutaric Aciduria Type 1 will result in inhibition of αKGDDs: this may be responsible for some of the clinical presentations seen in this disease.

In this study we show that the E2 subunit of the PDHc is a major target of glutarylation in CD8^+^ T cells. Glutarate in its CoA form, glutaryl-CoA, acts as a substrate for the recently discovered PTM glutarylation (Tan *et al*., 2014). We provide evidence that lysine glutarylation in the E2 subunit of the PDHc disrupts normal lipoyl function, resulting in disturbed PDHc activity; this is correlated with increased pyruvate conversion to lactate and increased glycolysis. This reaction occurs rapidly in cells treated with glutarate.

PDHc phosphorylation is also mediated by glutarate inhibition of HIF-P4H, and by the subsequent transcriptional induction of PDHK1; we have only seen this occur in cells cultured for long periods in DEG. Thus, acute changes in PDH activity and the subsequent effects on glycolysis observed here are most likely due to glutarylation of E2, whilst effects seen during long term culture in DEG are likely a more complex result of an interplay between glutarylation of PDHE2 and phosphorylation of PHDc by PDHK1. Regardless, glutarate is a potent and novel regulator of PDHc activity and clearly alters cellular metabolism via this interaction. It has been recently shown that PDHc inhibition in CD8^+^ T cells increases cytotoxicity and reduces tumor growth *in vivo* (Elia *et al*., 2022)which supports our observation here that glutarate can act to inhibit PDHc and can alter CD8^+^ T cell function in part via this inhibition.

An important aspect of these studies is our finding that glutarate levels change significantly in CD8^+^ T cells following T cell receptor activation. We also see that CD8^+^ T cell glutarate levels are dependent on both activation status and oxygen availability. Whilst most *in vitro* studies are carried out at 21% atmospheric oxygen levels, oxygen levels in lymphoid organs range from 0.5-4.5% (Caldwell *et al*., 2001), and when recruited to areas of insult, such as the tumor microenvironment, CD8^+^ T cells encounter even lower oxygen levels (as little as 0.1%). CD8^+^ T cells must therefore be flexible in their ability to function in different oxygen tensions and it has been shown that one way in which they orchestrate adaptation is altering cellular metabolic profiles (Doedens *et al*., 2013; Tyrakis *et al*., 2016). While it is established that the metabolites lactate and S-2HG are increased in CD8^+^ T cells in hypoxia (Tyrakis *et al*., 2016), glutarate had not been shown to be a hypoxia induced metabolite prior to this study. Similarly, after activation CD8^+^ T cells undergo massive metabolic changes to facilitate their increased energy demand needed to support proliferation and effector function (Ma *et al*., 2019). The activation induced increases in glutarate shown here further highlight the importance of this metabolite in T cell function.

Again, while we employed CD8+ T cells as our model system for this study, we show that glutarate induced inhibition of aKGDDs, and E2 glutarylation and subsequent regulation of PDHc activity are not restricted to T cells. This is important to consider given that glutarate has been found in recent metabolomics studies to be increased in physiological settings, e.g., post-exercise (Sato *et al*., 2022).

In this study we reveal a potentially central role for glutarate as a modulator and regulator of CD8^+^ T cell metabolism and cytotoxicity. We illustrate that glutarate can influence CD8^+^ T cell differentiation and increase effector function. We show that glutarate is an inhibitor of αKGDDs, which has many implications for better understanding the epigenetic and transcriptional landscape of cells. We additionally show that PDHE2 is a direct target of glutarate mediated glutarylation and is involved in the control of this crucial metabolic complex. Finally, we highlight the treatment potential of glutarate in an immunotherapeutic model, illustrating the translational potential for these findings.

## Acknowledgements

In-gel digestion, peptide extraction, mass spectrometric analysis and database searches for protein identification were carried out with thanks at the Proteomics Biomedicum, Karolinska Institute, Stockholm (https://ki.se/en/mbb/proteomics-biomedicum). We thank the animal staff at the Gurdon Institute Cambridge and Karolinska Institute for animal and technical assistance. We thank the flow cytometry facility from the School of the Biological Sciences Cambridge for their support and assistance in this work. This work was funded by Wellcome Trust Senior Investigator award to RSJ; the Evelyn Trust Cambridge (Patrick Sisson’s Research Fellowship) and the Karolinska Institute (Jonas Söderquists Fellowship) awarded to IPF; the Foundation for Science and Technology scholarship (SFRH/BD/115612/2016) awarded to PPC and Canadian Institutes of Health Research Fellowship to BJW; Wellcome Senior Fellowship to JAN (215477/Z/19/Z).

## Author Contributions

Conceptualization, E.M., P.P.C., R.S.J; Methodology, E.M. and P.P.C.; Investigation, E.M., P.P.C., A.Q., J.Z., S.S.T., B.W., R.H., G.G., and L.B.; Writing – Original Draft, E.M.; Writing – Review & Editing, E.M., P.P.C., P.V., D.B., C.W., J.A.N., P.K., I.P.F., and R.S.J.; Project Administration, R.S.J.; Funding Acquisition, R.S.J.

## Materials and Methods

### Animals

C57BL/6J animals (632, Charles River), were used in *in vitro* assays and in orthotopic tumor growth and infiltration experiments, in accordance with the ethical regulation of the UK home Office and the University of Cambridge. C57BL/6J (CD45.2) animals were purchased from Janvier Labs. Donor TCR-transgenic OT-1 mice (003831, The Jackson Laboratory) were crossed with mice bearing the CD45.1 congenic marker (002014, The Jackson Laboratory). Targeted deletion of HIF1 in T cells was achieved by crossing homozygous mice carrying loxP-flanked alleles Hif1 (Ryan *et al*., 1998) into a mouse strain of cre recombinase expression driven by the distal promoter of the lymphocyte-specific Lck gene (012837, The Jackson Laboratory). All the experiments were performed with age and sex matched cre negative controls. TCR-transgenic OT-1 mice and mice with targeted deletion of HIF1 in T cells were bred and housed in specific pathogen-free conditions in accordance with the regional animal ethics Committee of Northern Stockholm, Sweden.

### Cell Lines

B16-F10 cells were originally purchased from ATCC (CRL-6475) and genetically modified to express ovalbumin, eGFP and neomycin phosphotransferase (Velica et al. 2021). The resulting ovalbumin expression B16F10 cells were cultured in DMEM high glucose with pyruvate (11995065 Thermo Scientific) containing 0.75 mg/mL G418 sulfate (10131027, Thermo Scientific). HEK293 cells were a gift from Prof. Dantuma (Karolinska Institute, Stockholm) and cultured in DMEM high glucose with pyruvate. SKOV3 was purchased from ATCC (HTB-77) and cultured in McCoy’s 5A Medium (16600082, Thermo Scientific). Raji-GFP-Luc cells were purchased from Biocytogen (B-HCL-010) and cultured in complete RPMI (21875, Thermo Fisher). Jurkat cells purchased from ATCC (TIB-152) and cultured in complete RPMI (21875, Thermo Fisher). RAW 264.7 cells were purchased from ATCC (TIB-71) and cultured in RPMI (21875, Thermo Fisher). Mouse embryonic fibroblasts (MEFs) were originally derived as described previously (Sim *et al*., 2018). Cells were cultured in Dulbecco’s modified eagle medium (DMEM). All media was supplemented with 10% fetal bovine serum (FBS) (10270106, Gibco), 100 units/mL penicillin and 100 μg/mL streptomycin (P4333, Sigma). All cells were cultured in incubators with 5% CO2 and at either 21% or 1% oxygen, as indicated. Cell number and viability was determined using ADAM^TM^-MC Automated Cell Counter (NanoEntek), CellDrop™ Automated Cell Counter (DeNovix) or TC20 Automated Cell Counter (BioRad).

### T-cell isolation and activation

Human peripheral blood mononuclear cells (PBMCs) were obtained from National Health Service (NHS) Blood and Transplant (NHSBT: Addenbrooke’s Hospital, Cambridge, United Kingdom) or Karolinska Hospital Service, Sweden. Ethical approval was obtained from the East of England-Cambridge Central Research Ethics Committee (06/Q0108/281) and consent was obtained from all subjects. PBMCs from healthy donors were isolated from blood using discontinuous plasma-Percoll gradients. Naïve CD8^+^ T cells were isolated using the Naïve CD8^+^ T Cell Isolation Kit (130-093-244, Miltenyi), in accordance to manufacturer instructions. Total CD8^+^ T cells were purified by either positive or negative magnetic bead sorting (130-045-201, 130-096-495, Miltenyi). All human CD8^+^ T cells were activated with anti-human CD3/CD28 dynabeads (11131D, Thermo Scientific) at a 1:1 cell-to-bead ratio and cultured in complete RPMI supplemented with 10% FBS, 100 units/mL penicillin, 100 μg/mL streptomycin and 30U/mL IL-2 (11147528001, Sigma).

Mouse CD8^+^ T cells were isolated from mouse spleens by positive selection. Incubation with Microbeads conjugated to monoclonal anti-mouse CD8α (Ly-2; isotype: rat IgG2a) antibody (Miltenyi, 130-049-401) was followed by magnetic bead isolation on a MACS column. WT mouse CD8^+^ T cells were activated with anti-mouse CD3/CD28 dynabeads (11453D, Thermo Scientific) at a 1:1 cell-to-bead ratio. Purified OT-1 CD8^+^ T cells were activated with 1 μg/mL of the OVA-derived peptide SIINFEKL (ProImmune). All mouse CD8^+^ T cells were cultured in complete RPMI supplemented with 10% FBS, 100 units/mL penicillin, 100 μg/mL streptomycin, 55 μM 2-ME (21985023, Gibco) and 10U/mL IL-2 (11147528001, Sigma).

### Flow Cytometry and Sorting

Single cell suspensions were stained with Near-IR Dead Cell Stain Kit (10119, Thermo Fisher) followed by surface and intracellular staining with fluorochrome-labelled antibodies (Supplementary Table 1). Staining of cytoplasmic and nuclear antigens was performed using the Fixation/Permeabilization kit (554714, BD Biosciences) and the Transcription Factor buffer set (562725, BD Biosciences), respectively. For proliferation assays, cells were loaded with Cell Trace Violet (C34557, Thermo Fisher) according to manufacturer’s instructions. Samples were acquired on an Aurora (Cytek Biosciences).

For 5hmC staining, cells were stained for surface antigens as above and then fixed and permeabilised with the Transcription Factor buffer set (562725, BD Biosciences). Next cells were incubated with 4M HCL for 10 min at room temperature. The cells were then thoroughly washed and incubated in blocking buffer (0.1% PBS-Triton, 5% FBS) for 30min at 4°C. The cells were then incubated with primary anti-5hmC (10013602; Active Motif) overnight at 4°C and the day after with secondary antibody for 1 hr at room temperature. Flow cytometry was then performed as explained above.

Cells were sorted on an Aria III (BD Biosciences) following surface antigen staining as described above.

Human Antibodies
|  |  |  |  |  |
| --- | --- | --- | --- | --- |
| CCR7 | PE | 3D12 | BD | 552176 |
| CCR7 | PE/Cy7 | 3D12 | BD | 557648 |
| CD127 | PE | A019D5 | Biolegend | 351303 |
| CD127 | APC | A019D5 | Biolegend | 351342 |
| CD137 | PE | 4B4-1 | Biolegend | 309804 |
| CD19 | PE | 4G7 | Biolegend | 392505 |
| CD19 | APC | 4G7 | Biolegend | 392504 |
| CD25 | BV510 | BC96 | Biolegend | 302639 |
| CD27 | PerCP/Cy5.5 | M-T271 | Biolegend | 356408 |
| CD3 | BV510 | HIT3a | BD | 564713 |
| CD34 (RQR8) | AF488 | QBEnd10 | R&D | FAB7227G |
| CD44 | FITC | BJ18 | Biolegend | 338803 |
| CD45RA | BV650 | HI101 | Biolegend | 304135 |
| CD45RO | BUV496 | UCHL1 | BD | 749888 |
| CD45RO | BV605 | UCHL1 | Biolegend | 304238 |
| CD62L | AF488 | DREG-56 | Biolegend | 304816 |
| CD62L | PerCP/Cy5.5 | DREG-56 | Biolegend | 304824 |
| CD8a | BUV395 | RPA-T8 | BD | 563796 |
| CD8a | AF700 | HIT8a | Biolegend | 300920 |
| CD95 | PerCP/Cy5.5 | DX2 | Biolegend | 305630 |
| GzmB | PerCP/Cy5.5 | QA16A02 | Biolegend | 372211 |
| GzmB | AF647 | GB11 | Biolegend | 515405 |
| LAG3 (CD223) | AF647 | 11C3C65 | Biolegend | 369304 |
| Perforin | Pacific Blue | dG9 | Biolegend | 308117 |
| Perforin | PerCP/Cy5.5 | dG9 | Biolegend | 308114 |
| TBET | PerCP/Cy5.5 | 4B10 | Biolegend | 644805 |
| TBET | PE/Dazzle594 | 4B10 | Biolegend | 644828 |
| TCF1 | PE | S33-966 | BD | 564217 |
| TIGIT (VSTM3) | PE/Cy7 | A15153G | Biolegend | 372713 |
| TIM3 (CD366) | BV605 | F38-2E2 | Biolegend | 345017 |
| TOX | eFluor660 | TXRX10 | eBiosciences | 50-6502-82 |

Mouse Antibodies
|  |  |  |  |  |
| --- | --- | --- | --- | --- |
| CD11b | AF647 | M1/70 | Biolegend | 101220 |
| CD11b | AF488 | M1/70 | eBiosciences | 53-0112-80 |
| CD11c | PerCP/Cy5.5 | N418 | eBiosciences | 45-0114-80 |
| CD127 | AF647 | A7R34 | Biolegend | 135020 |
| CD137 | PE | 17B5 | Biolegend | 106105 |
| CD137 | APC | 17B5 | Biolegend | 106110 |
| CD19 | FITC | eBio1D3 | eBiosciences | 11-0193-82 |
| CD19 | PE | eBio1D3 | eBiosciences | 17-0193-82 |
| CD25 | PE | PC61.5 | eBiosciences | 12-0261-818 |
| CD27 | AF700 | LG.3A10 | Biolegend | 124239 |
| CD34 | AF647 | HM34 | Biolegend | 128605 |
| CD4 | BV421 | GK1.5 | Biolegend | 100437 |
| CD4 | AF488 | GK1.5 | Biolegend | 100423 |
| CD44 | PerCP/Cy5.5 | IM7 | Biolegend | 103032 |
| CD45.1 | PerCP/Cy5.5 | A20 | eBiosciences | 45-0453-82 |
| CD45.1 | PE | A20 | Biolegend | 110708 |
| CD45.2 | AF488 | 104 | Biolegend | 109815 |
| CD62L | AF488 | MEL-14 | Biolegend | 104419 |
| CD62L | PE | MEL-14 | Biolegend | 104408 |
| CD8a | BUV395 | 53-6.7 | BD | 565968 |
| CD8a | AF700 | 53-6.7 | Biolegend | 100730 |
| CTLA4(CD152) | PerCP/Cy5.5 | UC10-4B9 | Biolegend | 106315 |
| GzmB | PerCP/Cy5.5 | AD2 | Biolegend | 344013 |
| LAG3(CD223) | PerCP/Cy5.5 | C9B7W | Biolegend | 125212 |
| PD-1(CD279) | FITC | 29F.1A12 | Biolegend | 135214 |
| Perforin | APC | eBioOMAK-D | Biolegend | 17-9392-80 |
| TBET | PerCP/Cy5.5 | 4B10 | Biolegend | 644805 |
| TCF1 | PE | S33-966 | BD | 564217 |
| TIM3(CD366) | BV605 | RMT3-23 | Biolegend | 119721 |

### ELISA

Secreted IFNγ and TNFα levels of cultured cells were determined using an enzyme linked immunosorbent assays (ELISA). Assays were performed with uncoated ELISA kits (88-7316 and 88-7346-88, Thermo Scientific), according to manufacturer instructions. Absorbance was determined using a microplate reader (Sunrise, Tecan Austria GmbH) at a wavelength of 450 nm.

### Lentivirus transductions

For generation of lentiviral particles, 5×10^6^ HEK293 cells were plated in 15cm petri dishes and transfected the day after with 50 μL FuGENE (E2311, Promega), 10 μg CAR-encoding vectors and 3.3 μg of each 3rd generation lentivirus helper vectors (CART-027CL, Creative-Biolabs). Supernatant media containing lentiviral particles was harvested 48 hours after transfection and used fresh or stored at −80 °C. Lentiviral supernatants were spun onto nontreated wells plates coated with 30 μg/mL Retronectin reused up to three times (T100B, Takara), at 2000 x g for 2 hours at 32°C and replaced with activated human CD8^+^ T cells in fresh RPMI supplemented with 30 U/mL IL-2.

### *In vitro* cytotoxicity

10000 B16F10-OVA or SKOV3 cells were seeded per well in 96 well plates and co-cultured for a minimum of 14 hours with varying ratios of mouse CD8^+^ OT-1 or human CD8^+^RQR8^+^ HER2-CAR-T cells, respectively. T cells had been treated with or without DEG 7 days prior to assay. Wells were washed twice with PBS to remove T cells and the number of remaining target cells was determined by culturing with 10 μg/mL resazurin (R7017, Sigma) and measuring fluorescence signal in a plate reader. Cytotoxicity was calculated relative to wells with no T cells added. 10000 Raji cells were co-cultured for a minimum of 14 hours with CD19-CAR-T cells previously activated for 7 days and cultured with or without DEG. Cytotoxicity was assessed by flow cytometry by the ratio of live Raji cells to CountBright Absolute counting beads (C36950, Thermo Fisher). To determine specific cytotoxicity, data was normalised to the cytotoxicity of VC-transduced CD8 T cells of the respective donor.

### Vectors

DNA encoding a codon-optimized polycistronic peptide composed of RQR8 and anti-human HER-2 (clone 4D5) interspersed with picornavirus T2A and furin cleavage sequences was synthesized by GeneScript. RQR8, used as the transduction marker, is a chimeric surface protein composed with domains from CD34 (for detection and purification with clone QBEND/10), CD8 (for anchoring at the cell surface) and CD20 (for depletion *in vivo* with anti-CD20 mAb rituximab) (Philip *et al*., 2014). MicroRNA embedded shRNAs were generated as previously described (Ros and Gu, 2016). Briefly, 97-mer oligonucleotides (IDT Ultramers) coding for the respective shRNAs (Moffat *et al*., 2006) were PCR amplified using 10 μM of the primers miRE-XhoI-fw (5’-TGAACTCGAGAAGGTATATTGCTGTTGACAGTGAGCG-3’) and miRE-EcoRI-rev (5’-TCTCGAATTCTAGCCCCTTGAAGTCCGAGGCAGTAGGC-3’), 0.5 ng oligonucleotide template, and the Q5 High-Fidelity 2X Master Mix (NEB), and cloned HER-2 CAR vectors containing the miRE scaffold sequence. All coding sequences were cloned into pCDCAR1 (Creative-Biolabs). Third generation lentiviral transfer helper plasmids were obtained from Biocytogen. The following lentiviral vectors were purchased by Creative Biolabs were used: a truncated form of epidermal growth factor receptor (tEGFR; vector-control), anti-CD19-CAR with a 4-1BB endodomain. All sequences and accession numbers are available in Supplementary Material.

### Metabolite extraction and LC/MS analysis

Cells were counted to determine viable cell numbers. 0.5-2e6 viable cells were harvested, washed with cold PBS, and metabolic activity quenched by freezing samples in dry ice and ethanol, and stored at −80 °C. Metabolites were extracted by addition of 150 μl ice-cold 1:1 (vol/vol) methanol/water (containing 1mM N-Cyclohexyl-2-aminoethanesulfonic acid (CHES) as an internal standard) to the cell pellets, samples were transferred to a chilled microcentrifuge tube containing 150 μl chloroform and 500 μl methanol (750 μl total, in 3:1:1 vol/vol methanol/water/chloroform). Samples were sonicated in a water bath for 8 min at 4°C, and centrifuged (13,000 rpm) for 10 min at 4°C. The supernatant containing the extract was transferred to a new tube for evaporation using TurboVap LV (Biotage), resuspended in 3:3:1 (vol/vol/vol) methanol/water/chloroform (350 μl total) to phase separate polar metabolites from apolar metabolites, and centrifuged. The aqueous phase was transferred to a new tube for evaporation using TurboVap LV (Biotage), washed with 60 μl methanol and dried again. Evaporated extracts were dissolved in 100ul MeOH - 50% water, prior to injection on the machine. Isotopically labelled glutarates were determined by LC-MS/MS on a Waters Acquity UPLC system coupled to a Xevo-TQ-XS mass spectrometer. A SeQuant ZIC-HILIC column (2.1 mm×100 mm, 3.5 μm, Merck, Darmstad, Germany) equipped with a guard column (ZIC-HILIC Guard, 20 × 2.1 mm) was used for the separation, with mobile phase A consisting of MilliQ water, 0.1% v/v formic acid and mobile phase B consisting of acetonitrile, 0.1% v/v formic acid. The column was thermostated at 40°C, the injection volume was set to 3 μL, and the gradient was carried out at a flow rate of 0.3 mL/min as follows: 5% A from 0-1 min, linearly increased to 30% from min. 1-5 and to 75% at min 5.5. The column was subsequently washed in 75% A for 2.5 min and re-equilibrated at initial conditions for 3 min. MS analyses were performed in MRM mode, with positive electrospray ionization (ESI) from minute 0 – 1.4 and negative ESI from minute 1.4 – 4.7. Positive ESI was conducted according to the following parameters: capillary potential 3 kV, source offset 30 V, cone voltage 15V, source temperature 150°C, desolvation temperature 600°C, cone gas flow 150 L/h, desolvation gas flow, 1000 L/h, nebulizer 7 bar, and collision gas flow 0.17 mL/min. Negative ESI was operated in the same conditions but with a capillary potential set to 2 kV and a cone voltage set to 20V. Acquisition was performed by multiple reaction monitoring (MRM) with the mass transitions and the parameters illustrated in Table X. Two transitions were used for each of the target compounds, and the ion ratio between them was used to increase the confidence in the identification.

### qPCR

Total RNA was extracted from isolated CD8^+^ T cells (RNeasy kit, Qiagen) and 300 ng of RNA were used for cDNA synthesis (First-Strand Synthesis kit, Invitrogen). Quantitative real-time PCR (qRT-PCR) was performed with SYBR green (Roche) in a StepOnePlus system (Applied Biosystems). All kits were used according to the manufacturer’s instructions. Samples were run in technical duplicates. Primers were designed with NCBI primer blast and are listed in Supplementary Table 3.

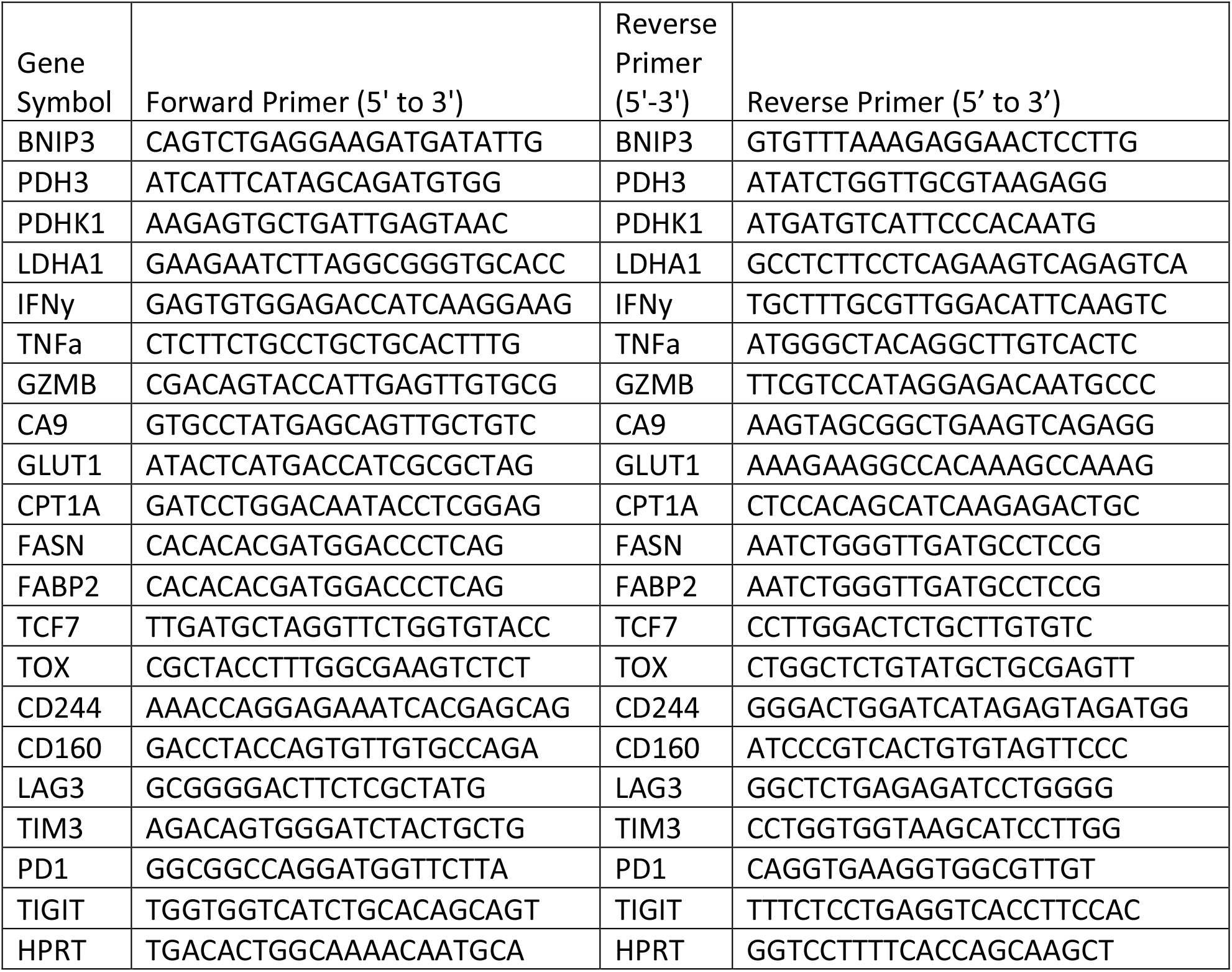

### Western Blotting

Cell pellets were lysed by either; urea-tris buffer (8 M urea, 50 mM Tris-HCl (pH=7.5), 150 mM β-mercaptoethanol), sonicated twice for 45 sec intercalated with 1 min incubation on ice and centrifuged at 14000 x g, 4 °C for 15 min or by RIPA Lysis and Extraction Buffer (89900, Thermo Scientific), according to manufacturer instructions. Nuclear and cytosolic fractions were prepared from cells with the NE-PER kit (78833, Thermo Scientific). Histones were extracted with Histone Extraction Kit (ab113476, ABCAM). Proteins were separated by SDS-PAGE and transferred to PVDF membranes. Total protein stain was obtained using non-stain protein labelling reagent (A44448, Thermo Scientific). Membranes were then blocked in 5% milk prepared in phosphate-buffered saline (PBS) plus 0.05% Tween 20, incubated with primary antibodies overnight at 4 °C and horseradish peroxidase (HRP)-conjugated secondary antibodies (HAF008 and HAF007, R&D) for 1 h the next day. Following ECL exposure (GERPN2106, Sigma), membranes were imaged using an iBrightCL1000 (Thermo Fisher). The following primary antibodies from cell signalling were used at a concentration of 1:1000: anti-PDHK1 (3820), anti-PPIB (43603), anti-Phospho-PDH a1 (37115), anti-SIRT5 (8779S), anti-H3 (4499), anti-H3K4me2 (9725), anti-H3K4me3 (9751), anti-H3K9me2 (4658), anti-H3K9me3 (13969), anti-H3K27me2 (9728), anti-H3K27me3 (9733), anti-H3K36me2 (2901), anti-H3K36me3 (4909), anti-H3K79me2 (5427), anti-H3K79me3 (4360), anti-H3k9ac (9649), anti-H3K18ac (13998) and anti-H3k27ac (8173). The following antibodies from Thermo Scientific were used at a concentration of 1:1000: PDHc (456799, Thermo Scientific), GCDH (PA5-60294). Other antibodies used at a concentration of 1:1000 are as follows: anti-HIF-1a (610958, BD), anti-HIF-1a (NB100-122, Novus), pan anti-glutarylysine (PTM-1151, PTM Biolabs) and pan anti-acetyllysine (PTM-105, PTM Biolabs).

### *In vitro* TET2 enzymatic activity assay

TET2 *in vitro* enzymatic activity assay was performed by Wuxi AppTec. the human TET2 enzyme (2 nM, BPS Bioscience) was incubated with the substrate of ssDNA (ssBiotin 26nt Me-C Oligo 30 nM, Genscript), in the presence of αKG (115 μM, Sigma), ammonium iron(II) sulfate hexahydrate (10 μM, Sigma), in assay buffer (50 mM HEPES pH 7.0, 100 mM NaCl, 0.01% Pluronic F-127, 1mM TCEP, 2mM ascorbic acid, 0.2 mg/ml BSA and 1000U/ml Catalase). DEG and glutaric acid were used in 3-fold serial dilution and the maximum concentration used was 5 mM. The pre-incubation time of the inhibitors with the TET2 mixture was 30 min at room temperature. The reaction step was at room temperature for 90 min. The product was detected by using an anti-5-Hydroxymethylcytosine antibody (5 nM, Active Motif), Eu-Protein A (5 nM, Cisbio), Streptavidin-Alexa Fluor 647 (6.25 nM, Life Technologies) and 10 mM EDTA (Sigma). For the standard curve, the ssBiotin 26nt HydMe-C Oligo (Genscript) was used. The assays were performed in technical duplicates in a 384-well plate. The inhibitor analysis was performed within the linear range of catalysis.

The percentage of inhibition was calculated with the following formula: Inhibition%=(1-(signal value per well-Average Low control)/(Average High control-Average Low control))*100. The data were fitted by Prism Graphpad with four parameters equation via “log(inhibitor) vs. response -- Variable slope” model.

### *In vitro* HIF-P4H-1 and KDM6A activity assays

The human full-length HIF-P4H-1 and KDM6A enzymes were recombinantly produced using Sf21 insect cells and the enzymes were purified with anti-FLAG M2 affinity gel as previously described (Chakraborty *et al*., 2019; Hirsilä *et al*., 2003). The proteins were analyzed by SDS-PAGE followed by Coomassie blue staining. The inhibition assays were carried out by a method which measures the hydroxylation-coupled stoichiometric release of ^14^CO_2_ from 2-oxo-[1-14C] glutarate with a synthetic HIF-1α peptides and a histone H3K27me3 peptide as substrates for HIF-P4H-1 and KDM6A respectively (Chakraborty *et al*., 2019; Hirsilä *et al*., 2003). Glutarate was used at increasing concentrations (50 μM to 10 mM) in the assay. The concentration of 2-oxoglutarate was 8 μM and 32μM for HIF-P4H-1 and KDM6A catalysed reactions, respectively, while keeping the concentrations of other cosubstrates and cofactors saturating and constant. The IC50 value of glutarate analogue was calculated from the inhibition saturation curves.

### HIF-PH cellular activity assays

MEFs were transiently transfected using Lipofectamine 2000 (Thermo Fisher) with 20 ng per expression plasmid for 2 x 104 cells per well in a 96-well plate. Cells were treated with transfection mixes for 8 hours.

Generation of plasmids for HIF-PH activity luciferase reporter assay has been previously described: luciferase reporter driven by Gal4 response element (GRE-luc) pFLAG-Gal4-mHIF1α NTAD ((Pereira *et al*., 2003). Briefly, cells were concurrently transfected with pGRE-luc and a plasmid encoding constitutive expression of a fusion protein linking a FLAG-tagged Gal4 DNA-binding domain with one of the transcription activation domains of murine hypoxia inducible factor 1a (HIF-1α). The N-terminal transcription activation domain (NTAD) contains proline residues that are hydroxylated by HIF-PH enzymes to target the protein for degradation, thus luciferase expression is controlled by HIF-PH hydroxylation activity against the pFLAG-Gal4-mHIF-1α NTAD. Following 8 hours of transient transfection with expression plasmids, cells were treated with experimental media containing metabolites and drugs as described in figure legends for 16 hours and then luciferase signal assessed using the Steady-Glo luciferase assay system (Promega) detected using a luminometer. Luciferase signal was normalised to that induced by 2.5 mM DMOG.

### Immunoprecipitation and protein isolation for mass spectrometry

10-30×106 cells were lysed with Pierce IP Lysis Buffer (87787, Thermo Scientific). Immunoprecipitation was performed using protein A dynabeads (10002D, Thermo Scientific) according to manufacturer instructions. 1-10ug of target antibody was used per reaction. Eluted proteins were separated by SDS-PAGE, gel was stained with InstantBlue Coomassie Protein Stain (ab119211, ABCAM), imaged using an iBrightCL1000 (Thermo Fisher) and stained target bands were cut and stored in water until analysis. Non-cut gels were transferred to PVDF membranes and then blocked in 5% milk prepared in phosphate-buffered saline (PBS) plus 0.05% Tween 20. Membranes were then incubated with primary antibodies overnight at 4°C (supplementary table 2) and horseradish peroxidase (HRP)-conjugated secondary antibodies for 1 h the next day. Following ECL exposure (GERPN2106, Sigma), membranes were imaged using an iBrightCL1000 (Thermo Fisher).

### In-gel Protein digestion and mass spectrometry

Protein bands were excised manually from gels and in-gel digested using a MassPREP robotic protein-handling system (Waters, Millford, MA, USA). Gel pieces were distained following the manufacturer’s description. Proteins then were reduced with 10 mM DTT in 100 mM Ambic for 30 min at 40oC and alkylated with 55 mM iodoacetamide in 100 mM Ambic for 20 min at 4°C followed by digestion with 0.3 mg trypsin (sequence grade, Promega, Madison, WI) in 50 mM Ambic for 5 h at 4°C. The tryptic peptides were extracted with 1% formic acid in 2% acetonitrile, followed by 50% acetonitrile twice. The liquid was evaporated to dryness on a vacuum concentrator (Eppendorf).

The reconstituted peptides in solvent A (2% acetonitrile, 0.1% formic acid) were separated on a 50 cm long EASY-spray column (Thermo Fished Scientific) connected to an Ultimate-3000 nano-LC system (Thermo Fisher Scientific) using a 60 min gradient from 4-26% of solvent B (98% acetonitrile, 0.1% formic acid) in 55 min and up to 95% of solvent B in 5 min at a flow rate of 300 nL/min. Mass spectra were acquired on a Q Exactive HF hybrid Orbitrap mass spectrometer (Thermo Fisher Scientific) in m/z 375 to 1500 at resolution of R=120,000 (at m/z 200) for full mass, followed by data-dependent HCD fragmentations from 17 most intense precursor ions with a charge state 2+ to 7+. The tandem mass spectra were acquired with a resolution of R=30,000, targeting 2×10^5^ ions, setting isolation width to m/z 1.4 and normalized collision energy to 28%.

Acquired raw data files were analyzed using the Mascot Server v.2.5.1 (Matrix Science Ltd., UK) and searched against SwissProt protein databases (20,368 human entries). Maximum of two missed cleavage sites were allowed for trypsin, while setting the precursor and the fragment ion mass tolerance to 10 ppm and 0.02 Da, respectively. Dynamic modifications of oxidation on methionine, deamidation of asparagine and glutamine and acetylation of N-termini were set. Initial search results were filtered with 5% FDR using Percolator to recalculate Mascot scores. Protein identifications were accepted if they could be established at greater than 96.0% probability and contained at least 2 identified peptides. Proteins that contained similar peptides and could not be differentiated based on MS/MS analysis alone were grouped to satisfy the principles of parsimony.

### PDHc activity assay

PDHc activity was determined using a PDHc Activity Assay Kit (MAK183, Sigma). 1×106 cells were used per reaction and assay was performed according to manufacturer instructions. Activity was normalised to total protein concentration.

### Seahorse

A Seahorse XFe bioanalyser was used to measure oxygen consumption rate (OCR) and extracellular acidification rate (ECAR). 1.5×105 CD8^+^ T cells per well were spun onto poly-D-lysine (P7280, Sigma) coated seahorse plates and preincubated at 37°C for a minimum of 30 min in the absence of CO2 in Seahorse XF RMPI medium, pH 7.4 (10.576-100, Agilent), supplemented with 10 mM glucose (A2494001, Thermo Scientific) and 2 mM glutamine (25030081, Thermo Scientific). A minimum of 5 technical replicates per biological replicate were used. For the mitochondrial stress test (MST) and glucose stress test (GST) OCR and ECAR were measured under basal conditions, and after the addition of the following drugs: 750 μM DEG (QB-1473, Combi-block), 0.05% DMSO, 1 μM oligomycin (75351, Sigma), 1.5 μM flurorcarbonyl cyanide phenylhydrazone (FCCP) (C2920, Sigma), 100 nM rotenone + 1μM antimycin A R8875; A8674, Sigma), 10mM glucose (A2494001, Thermo Scientific and 50mM 2-DG (B1048-100, BioVision) as indicated. To asses mitochondrial fuel usage OCR was measured subsequent to the addition of the following drugs in different combinations as indicated: 3 μM bis-2-(5-phenylacetamido-1,3,4-thiadiazol-2-yl)ethyl sulfide (BPTES) (SML0601, Sigma), 2 μM UK-5099 (PZ0160, Sigma) and 4 μM etomoxir (E1905). Assay parameters were as follows: 3 min mix, no wait, 3 min measurement, repeated 3–5 times at basal and after each addition. Measurements were taken using a 96 well Extracellular Flux Analyzer (Seahorse bioscience).

### Orthotopic tumor growth and infiltration experiments

7 to 10-weeks old male C57BL/6j mice were inoculated subcutaneously with 1.0×10^6^ B16F10. Animals were assigned randomly to each experimental group. 4 days post inoculation, mice were treated with an intraperitoneal injection of 10mg/kg DEG. Treatment was repeated every 2-3 days until day 28. Tumor volume was measured every 2-3 days with electronic callipers until day 30. Tumor volume was calculated using the formula a×b×b/2 where a is the length and b is the width of the tumor. Peripheral blood was collected from the tail vein at days 7 and 14 days after tumor inoculation and analysed by flow cytometry. On day 14, tumors were dissected and dissociated using the Tumor Dissociation Kit (130-095-929, Miltenyi), according to manufacturer instructions. Spleen and the tumor draining lymph node were mashed in cell strainers. The tumor, spleen and draining lymph node single-cell suspensions were stained with fluorochrome-labelled antibodies and analysed by flow cytometry.

### Statistics

Statistical analyses were performed with Prism 9 software (GraphPad). A P value of <0.05 was considered significant and the statistical tests used are stated in figure legends.

## Supplemental Figures

**Supplemental Figure 1:**
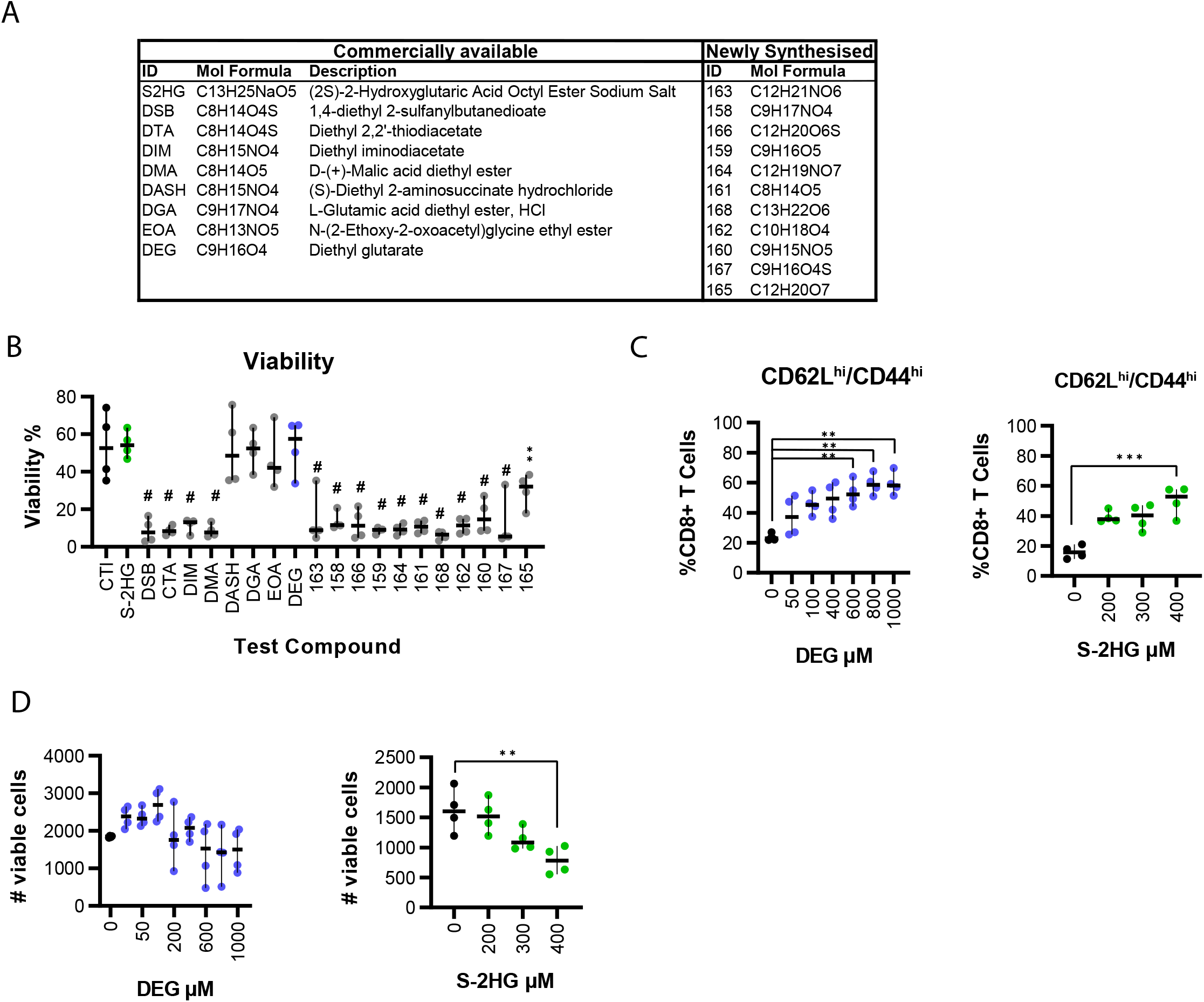
DEG is a modulator of T cell differentiation. (A) List of 19 test compounds (plus S-2HG) used in screen in described in Figure 1A. (B) Percentage viability of CD8^+^ T cells following 7 days of treatment with 400μM of test compound, as determined by flow cytometry. Ordinary one-way ANOVA relative to CT; n=4. (C) Percentage of CD62L^hi^/CD44^hi^ CD8^+^ T cells following 7 days of treatment with increasing concentrations of DEG (left) or S-2HG (right). Ordinary one-way ANOVA; n=4. (D) Number of viable CD8^+^ cells following 7 days of treatment with increasing concentration of DEG (right) or S-2HG (left), as determined by counting beads and flow cytometry. Ordinary one-way ANOVA relative to CT; n=4. All scatter plots show median and 95% CI where each dot represents one murine donor. *p<0.05, **p<0.01, ***p<0.001, #p<0.0001.

**Supplemental Figure 2:**
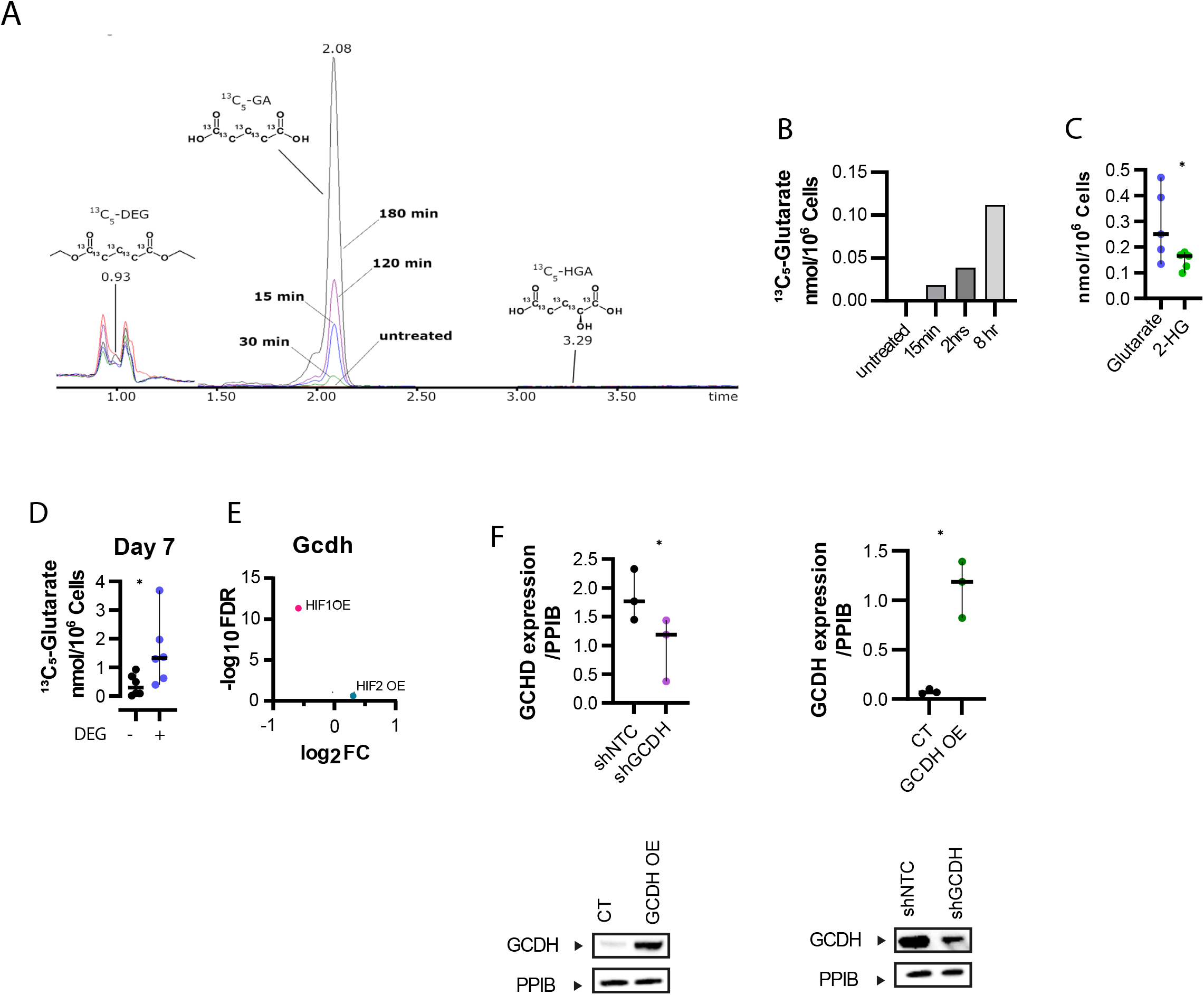
Glutarate is a novel T cell metabolite with immunomodulatory properties. (A) Chromatograms of isotope tracing. Human CD8^+^ T cells were treated with ^13^C5-DEG for varying time length and mass spectrometry for glutarate was performed. (B) Quantified ^13^C5-Glutarate from cells treated with ^13^C5-DEG for varying time lengths, as described and shown as chromatograms in Figure S2A. (C) Glutarate and 2HG levels in activated human CD8^+^ T cells. Wilcoxon test: n=5. (D) Glutarate levels in human CD8^+^ T cells following 7 days of treatment +/− DEG 500μM. Paired t test; n=6. (D) Glutarate levels in human CD8^+^ T cells following 7 days of treatment +/− 500μM DEG. Paired t test; n=6. (E) *Gcdh* levels in transcripts HIF1 OE (HIF1AAA) and HIF2 OE (HIF2AAA) – transduced mouse CD8^+^ T cells, relative to vector control (Veliça *et al*., 2021). (F) GCDH levels in T cells transduced with an shGCDH or GCDH overexpressing vector as determined by western blot analysis. Paired t test; n=3. All scatter plots show median and 95% CI where each dot represents one human donor. *p<0.05, **p<0.01, ***p<0.001, #p<0.0001.

**Supplemental Figure 3:**
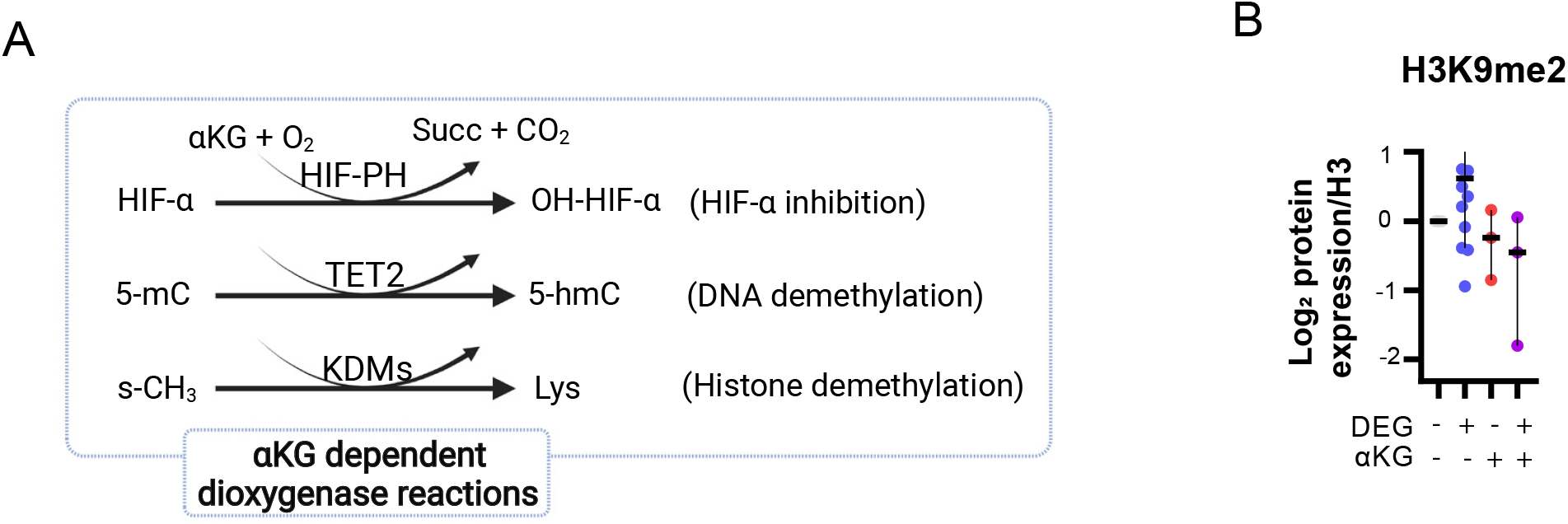
Glutarate is an inhibitor of αKGDDs. (A) Illustration of alphaketoglutarate (αKG) dependent dioxygenase (αKGDD) reactions investigated in this study. (B) Representative western blot and log2 FC protein expression of H3K9me2 in human CD8^+^ T cells treated with +/− DEG and/or αKG for 7 days. Graph shows median + 95% CI. Each dot represents one human donor. Kruskal-Wallis test; n=3-9.

**Supplemental Figure 4:**
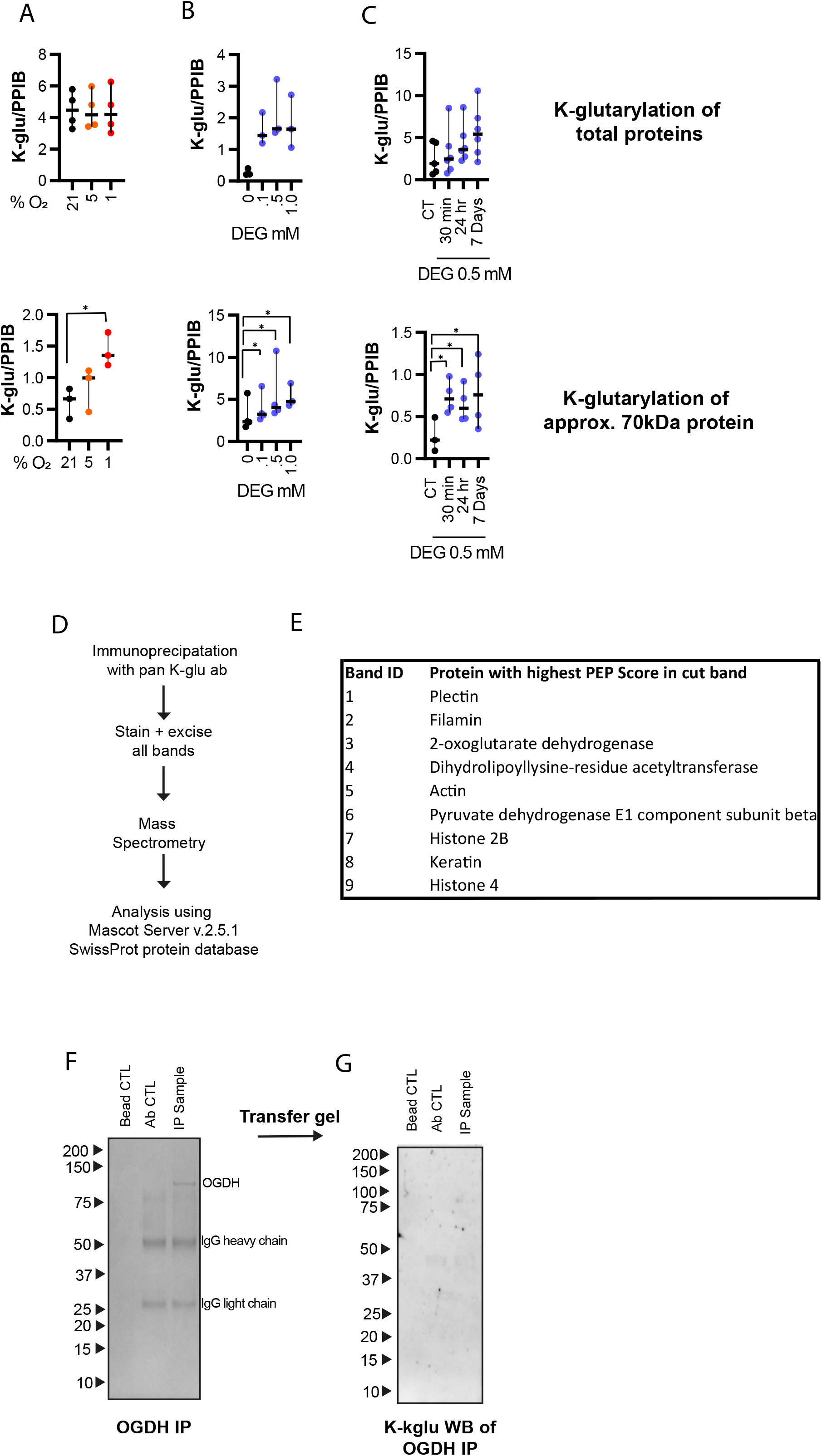
Glutarylation of PDHE2. (A) Quantified total protein lysine-glutarylation or lysine-glutarylation of 67-70kDa protein, as determined by western blot analysis and relative to PPIB, of cells treated with increasing concentrations of DEG for 7 days (as described in Figure 4D). Ordinary one-way ANOVA; n=3. (B) Quantified total protein lysine-glutarylation or lysine-glutarylation of 67-70kDa protein, as determined by western blot analysis and relative to PPIB, of human CD8^+^ T cells 7 days post activation. Cells were treated with DEG 500μM for different time lengths as indication and harvested at the same time 7 days post activation (as described in Figure 4E). Ordinary one-way ANOVA; n=3. (C) Quantified total protein lysine-glutarylation or lysine-glutarylation of 67-70kDa protein, as determined by western blot analysis and relative to PPIB, of human CD8^+^ T cells which were cultured at different oxygen tensions for 24hr prior to harvest (as in described in Figure 4C). (D) Model of lysine-glutarylated protein isolation and identification by mass spectrometry. Lysine-glutarylation immunoprecipitation was performed on 30×10^6^ mouse CD8^+^ T cells, 7 days post activation. Following protein separation by SDS-PAGE, gel was stained with Coomassie blue and stained bands were excised, gel was digested and mass spectrometry was performed. Data was analysed using the Mascot Server v.2.3.1 and the SwissProt protein database. 3 independent experiments and analysis performed. (E) Top hits for each excised band from mass spectrometry described in Figure 4G. (F) Oxoglutarate dehydrogenase (OGDH) complex isolated from 30×10^6^ mouse CD8^+^ T cells by immunoprecipitation. (G) k-glutarylation WB of immunoprecipitated OGDH complex as described in S4I. All scatter plots show median and 95% CI where each dot represents one human donor. *p<0.05.

**Supplemental Figure 5:**
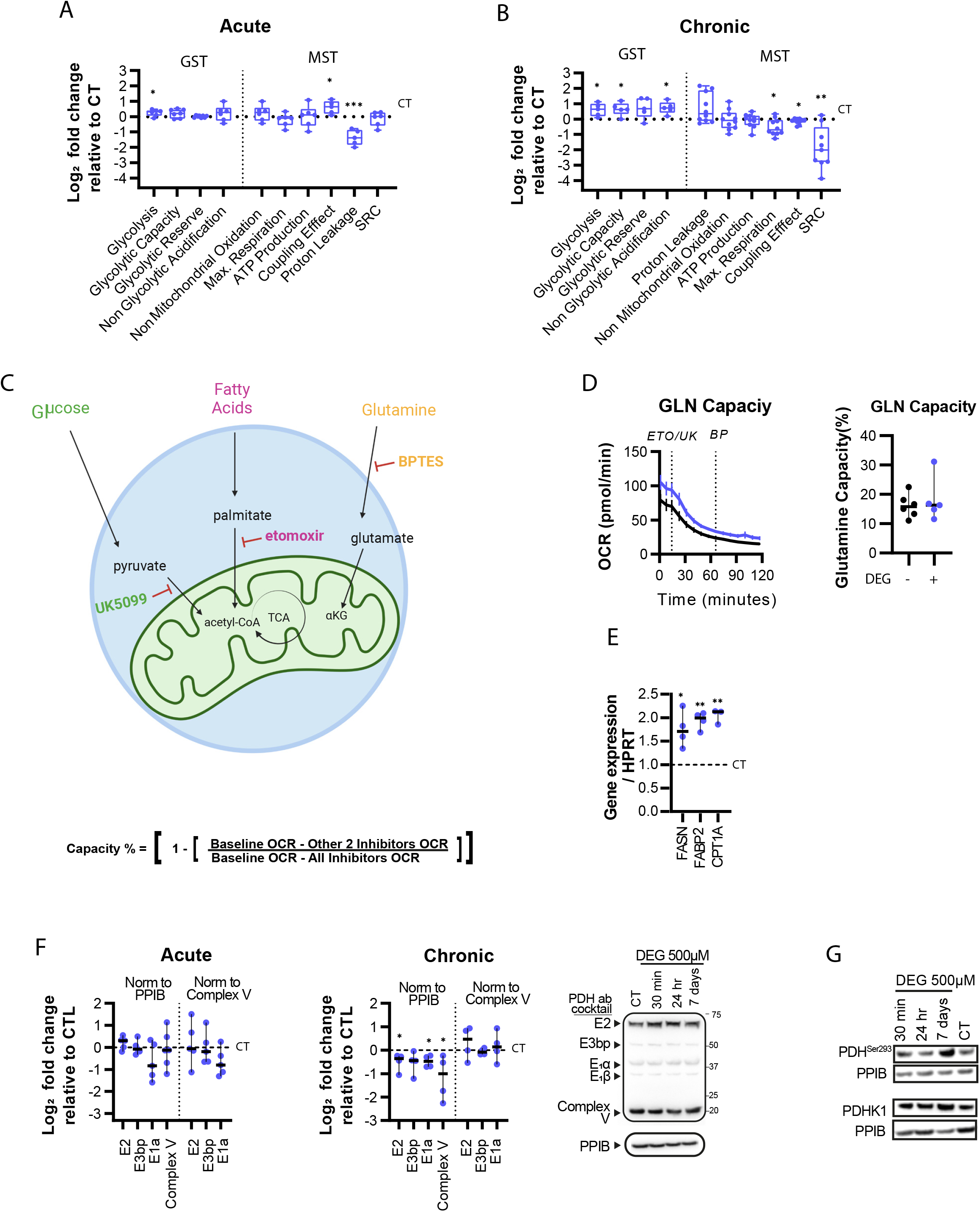
Glutarate modulates metabolism via both glutarylation and phosphorylation of the PDHc. (A and B) Quantitative analysis of MST and GST following acute (30 min) (left) and chronic (7 days) (right) DEG exposure, as described in Figure 5H-J, shown as log2 FC relative to untreated control (black dotted line). One sample t test; n=6-10. (C) Model of mitochondrial fuel flexibility tests. (D) Glutamine oxidative capacity of CD8^+^ T cells treated +/− DEG 500 μM for 7 days, as determined by OCR measurement pre and post addition of BPTES (BP), UK5099 (UK) and etomoxir (ETO) as indicated. Paired t test; n=5. (E) Gene expression of CD8^+^ T cells following 7 days of treatment with DEG, as determined by qPCR analysis, and represented as fold change relative to untreated control (dotted black line). One sample t test; n=3-4. (F) Protein expression of PDHc in CD8^+^ T cells treated +/− DEG 500μM. Acute treatment of CD8^+^ T cells 7 days post activation for 30 min. Chronic treatment of CD8^+^ T cells for 7 days, from activation. Protein levels normalised to PPIB or complex V expression as indicated. One-sample t-test; n=4. (G) Representative western blot of CD8^+^ T cells 7 days post activation as described and shown in Figure 5N. Cells were treated with DEG 500μM for different time lengths as indication and harvested at the same time 7 days post activation. All scatter plots show median and 95% CI where each dot represents one human donor. *p<0.05, **p<0.01, ***p<0.001.

**Supplemental Figure 6:**
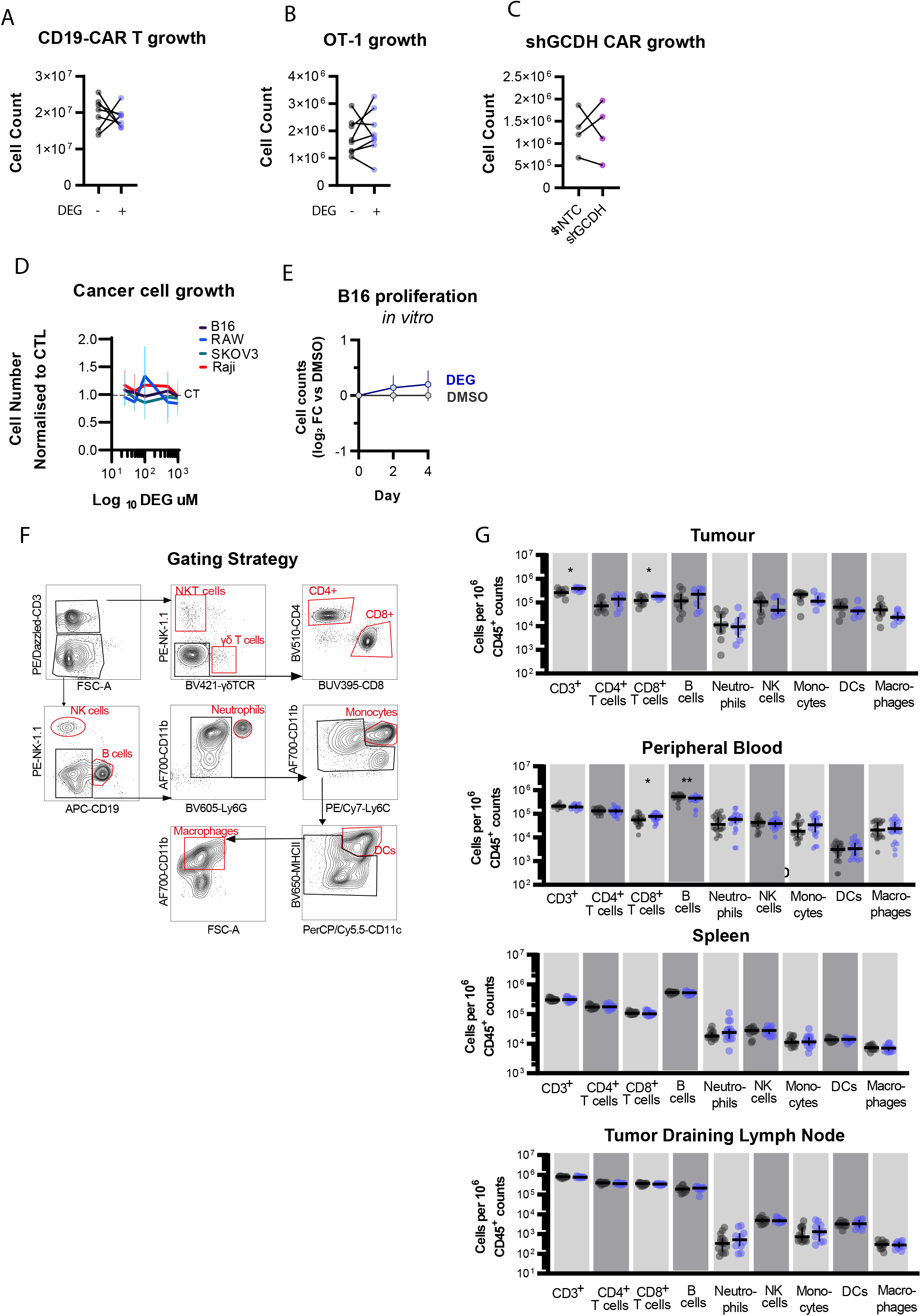
Glutarate reduces tumor growth, and increases T cell numbers and tumor infiltration. (A) Cell count of human CAR-T cells treated +/− 500μM for 7 days. Paired t test; n=6. (B) Cell count of murine OT-1 CD8^+^ T cells treated +/− 500μM for 7 days. Paired t test; n=6. (C) Cell count of HER2-CAR-T_shRNA cell numbers 7 days post transduction. Paired t test; n=4. (D) Cell count of different cancer cell lines treated with increasing concentrations of DEG for 72hr. Cell count relative to appropriate untreated control. One way ANOVA; n=3. (E) *In vitro* B16F10 cell growth +/− DEG 500μM as determined by cell counting and represented as log2 FC. One way ANOVA; n=3. (F) Gating strategy used for analysis of Figure 6K-L and Figure S6G. (G) Frequency of cell types in peripheral blood, tumor, spleen and tumour draining lymph nodes, 14 days post tumor inoculation, represented as log2 FC relative to CD45^+^ count. Graph shows median + 95% CI where each dot represents one mouse. Unpaired t test; n=14-16. *p<0.05, **p<0.01.

